# Fleming: An AI Agent for Antibiotic Design for Mycobacterium tuberculosis

**DOI:** 10.1101/2025.04.01.646719

**Authors:** Ziming Wei, Yasha Ektefaie, Andrew Zhou, Dereje Negatu, Bree Aldridge, Thomas Dick, Michael Skarlinski, Andrew D. White, Samuel Rodriques, Sepideh Hosseiniporgham, Maloy Parai, Armando Flores, Inna Krieger, Marinka Zitnik, James Sacchettini, Maha Farhat

## Abstract

Antibiotic development is challenged by high costs and failure rates. Artificial intelligence (AI) holds promise to overcome these challenges by predicting inhibitory properties of novel compounds, generating new candidates, and contextualizing property predictions in the biological background. Fleming is an integrative AI agent that explores novel chemical space to identify lead compounds meeting multiple criteria. The discriminative and generative AI models for Mycobacterium tuberculosis (Mtb) inhibition were trained on a set of 114,900 diverse compounds and fragments based on *in vitro* growth inhibition. We combined both models as well as molecular optimization, ADMET prediction and literature search functions to make Fleming an integrated agent for Mtb preclinical lead identification. Fleming has 17% higher discrimination between known Mtb leads and leads for other diseases than a generic LLM agent along with 13% higher discrimination than molecular property prediction alone on challenging ADMET tasks. Fleming demonstrates an 83% *in vitro* hit rate of predicted inhibition and a 100% hit rate of *de novo* generative design. Fleming’s generative designs also demonstrate an 83% rate of favorable ADMET profiles. Fleming is an integrative AI agent able to explore new regions of the chemical space to select lead compounds that simultaneously meet several desirable criteria.

## Background

Multidrug-resistant (MDR) strains of *Mycobacterium tuberculosis* (Mtb) caused approximately 160,000 deaths in 2022 and account for over 13% of global deaths attributable to antimicrobial resistance [1, 2]. Mathematical models further estimate that nearly 19 million individuals harbor latent MDR-TB infections and remain at risk of progression to active disease [1, 2]. The persistence of MDR-TB reflects the prolonged chemotherapy required for cure and continued reliance on antibiotics that have been in continuous use for more than five decades. Consequently, there is an immediate need for new tuberculosis (TB) antibiotics that target novel mechanisms and enable shorter, more effective treatment regimens [3].

High-throughput antibacterial screening (HTS) enables the evaluation of hundreds of thousands of compounds for whole-cell growth inhibition, but it samples a limited and highly redundant chemical space [4, 5]. Despite decades of effort, HTS-driven discovery of TB-active molecules with favorable properties has been slow, costly, and labor intensive[6]. These challenges underscore the necessity for alternative approaches capable of efficiently exploring underrepresented regions of chemical space while satisfying complex biological constraints.

Artificial intelligence (AI) has emerged as a powerful paradigm for accelerating drug discovery. In infectious diseases, AI offers particular promise by enabling rapid screening of vast chemical libraries, identification of novel molecular scaffolds capable of penetrating bacterial envelopes, and prediction of mechanisms of action that may elude conventional methods [7–9]. In principle, AI can also identify synergistic drug combinations and minimize cross-resistance with existing antibiotics. However, realizing this potential requires integration of predictive models for efficacy with constraints on absorption, distribution, metabolism, excretion, and toxicity (ADMET), chemical stability, and synthetic accessibility.

Numerous computational models have been developed to predict molecular properties from chemical descriptors and structural information. These range from traditional approaches, such as regression and random forests[10–17], to modern architectures including graph neural networks (GNNs) and transformers that directly model molecular structure [18–29]. Although these models often report high benchmark performance, their predictive accuracy can degrade substantially when applied to structurally novel compounds, and uncertainty estimates are frequently poorly calibrated [15, 30–32]. Generative models aim to address this limitation by designing molecules *de novo*, but current approaches—including Monte Carlo tree search and diffusion models—remain constrained by training data biases and have yet to yield viable drug candidates [33] [34].

Lead discovery ultimately requires the integration of diverse chemical and biological knowledge, traditionally distributed across multiple domain specialists. An effective AI platform must therefore unify predictive models across domains, account for model uncertainty, and incorporate external knowledge from medicinal chemistry literature and known structure–property relationships. Large language models (LLMs) provide a compelling framework for integrating heterogeneous textual and molecular information and for acting as agents that coordinate multiple predictive tools within an interactive, multi-objective discovery workflow [35][36].

Here, we present **Fleming**, an AI-driven, multi-domain platform designed to accelerate the discovery of next-generation TB antibiotics. Fleming integrates a state-of-the-art Mtb whole-cell inhibition GNN trained on over 114,000 compounds with generative agents for conditional *de novo* design, ADMET prediction models, molecular optimization tools, and advanced literature-driven reasoning. In prospective validation, Fleming demonstrates substantial enrichment over random screening and identifies compounds with favorable pharmacokinetic and toxicity profiles, clearly demostrating the power of integrative AI agents to navigate unexplored chemical space while satisfying complex biological requirements.

## Results

### Overview of Fleming

Fleming consists of a central medicinal chemist agent that orchestrates four sub-agents: the Mtb growth inhibition agent, the generative agent, the ADMET agent, and the molecular optimizer agent (Figure 1a). The sub-agents interact directly with 9 molecular AI models and 11 tools ranging from standard cheminformatics methods to custom utilities including a natural language molecular describer and a molecular novelty calculator (Figure 1b). Fleming can either analyze or generate a small number of molecules in real-time (co-pilot mode), or batch-process large libraries to identify and prioritize promising candidates (batch mode).

**Figure 1:**
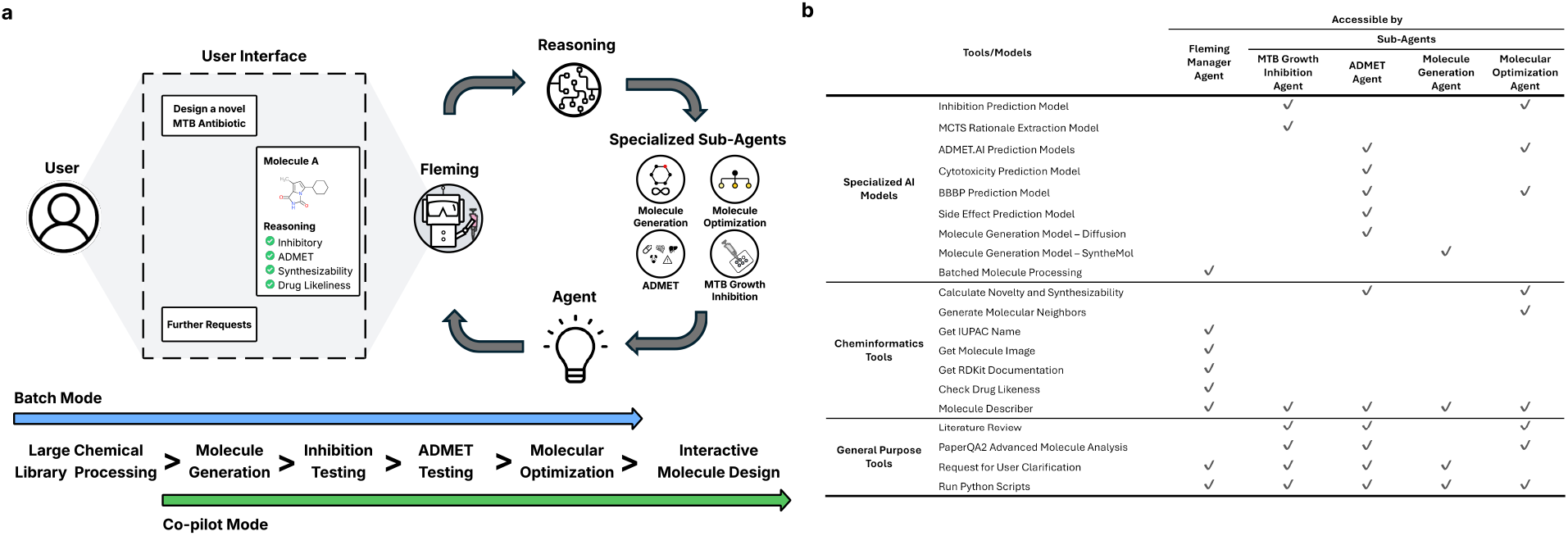
Fleming is an agent for antibiotic discovery in Tuberculosis (TB). **(a)** Fleming integrates four specialized agents — the ADMET agent, MTB growth inhibition agent, molecule generation agent, and molecular optimization agent — to perform tasks critical to designing novel antibiotics for MTB. **(b)** Fleming includes a diverse toolbox that includes machine learning models and chem-informatics tools. Different sub-agents have access to different sets of tools, which enables them to respond to user requests.

### Accurate Mtb inhibition predictions

We performed high-throughput screening of drug-like small molecules for inhibition of Mtb (Figure 2a). The set consisted of 113,919 compounds (molecular mass IQR 304-406 g/mol) assembled for high diversity by clustering the structures of drug-like molecules from the commercial chemical space and selecting the most representative molecules from each cluster. The resulting compound library shows substantial structural diversity, with no pair of molecules exhibiting a Tanimoto similarity greater than 0.65, and all compounds conform to a relaxed interpretation of Lipinski’s drug-likeness criteria [37] (Supplementary Methods). Compounds were tested at relatively low concentrations (10µM - 20µM), and viability of mc^2^-7000 Mtb strain (PMID: 12219086) [38] was assessed using two orthogonal signals - luminescence of luciferase expressed from a plasmid and resazurin reduction (PMID: 25779323) [39] - independently or in combination (Methods). The normalized percent inhibition readouts were used to assign molecules with 40% or greater growth inhibition as positives. We pooled this data with Mtb growth inhibition measurements of 1000 molecular fragments [40] (molecular mass IQR 154-203 g/mol, Methods) and 14 current TB antibiotics. The final training data contained 6,572 (5.7%) Mtb growth inhibitory compounds or fragments and 108,361 (94.3%) non-inhibitors (Figure 2a).

**Figure 2:**
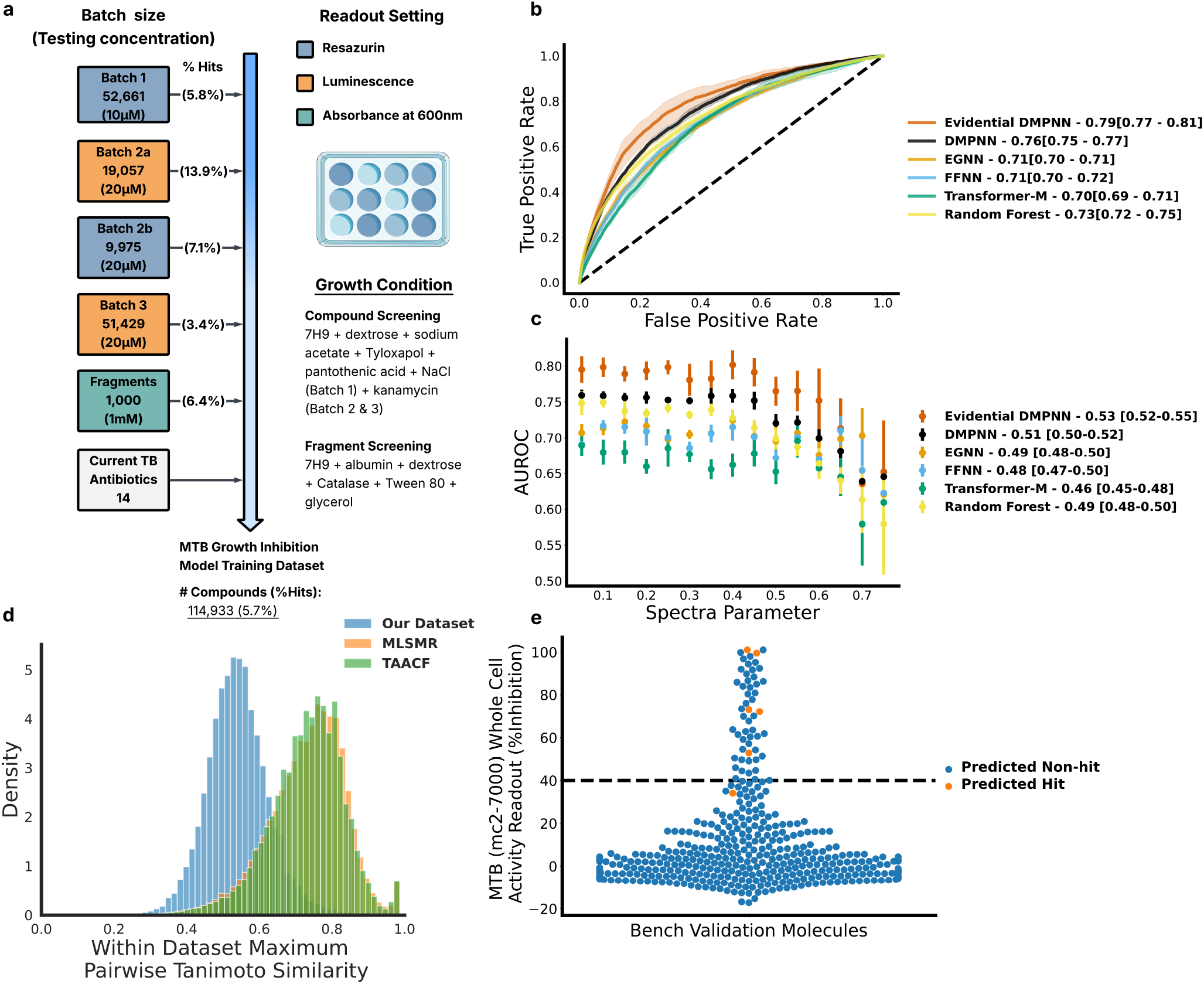
Inhibition agent identifies small molecules with inhibitory effects on TB. **(a)** A high-quality, diverse MTB growth inhibition dataset underpins the development of these agents, particularly the inhibition prediction and molecular generation modules. The dataset comprises 114,933 compounds and fragments, including three batches of high-throughput compound screening, one batch of fragment screening, and 14 known anti-TB antibiotics. The compound screening batches were conducted under slightly varied growth conditions and readout parameters to maximize chemical and phenotypic diversity. The overall hit rate of the dataset is 5.7%. **(b)** Area under receiver operating characteristic (AUROC) curve of benchmarked models on TB inhibition prediction. **(c)** Area under the spectral performance curve (AUSPC) of benchmarked models on TB inhibition prediction. AUSPC generated with spectral parameters from 0.00 to 0.75 with step size of 0.05. **(d)** Our training data show significantly (*p* − *value <* 0.0001) lower maximum pair-wise similarity within the dataset, which indicates that our dataset has a higher compound diversity that enables the training of a more generalizable model. **(e)** %inhibition readouts of testing set molecules on whole-cell activity against Mtb(mc27000). 40% is used as the threshold of inhibition binarization. Blue dots are predicted non-hits, and orange dots are predicted hits. Predicted hits are defined as the molecules with predicted inhibition *>* 0.7 and prediction confidence *>* 0.6.

Five advanced molecular property predictors were trained to estimate the inhibition across the 114,933 molecules. We compared a directed message passing neural network (DMPNN) [18], a specialized graph neural network (EGNN) [19], a molecular transformer integrating both 2D and 3D molecular representations (Transformer-M) [41], a feed-forward neural network, and random forests [42]. The DMPNN performs best on the two most widely used measures of binary classifiers: Area Under the Receiver Operating Curve - AUROC = 0.76, and Area Under the Precision-Recall Curve - AUPRC = 0.18; paired t-test, *p <* 0.001 five-fold cross-validation. Model performance was evaluated using the area under the receiver operating characteristic curve (AUROC), which quantifies the ability of the model to rank true positive instances higher than true negatives across decision thresholds, and the area under the precision-recall curve (AUPRC), which summarizes the model’s capacity to identify true positives while limiting false positive predictions. The DMPNN was also the most generalizable property prediction model on test molecules with decreasing similarity to the training molecules (using SPECTRA [43], Area Under the Spectral Parameter Curve - AUSPC = 0.35) (Methods; Figure 2b,c). We trained the final DMPNN using the evidential framework (eDMPNN) by adding a Dirichlet loss [44] to capture uncertainty in inhibition model prediction from noisy training data or out-of-distribution effects (Methods). Uncertainty filtering resulted in significant improvement in performance with the AUROC increasing from 0.76 to 0.79, (paired t-test p-value = 0.02), and the AUPRC from 0.18 to 0.20 (p-value = 0.01). AUSPC from 0.51 to 0.53. (Figure 2c,d, Supplementary Figure 1c).

External retrospective validation was based on two references: an Mtb HTS dataset (214,507 compounds from the Molecular Libraries Small Molecule Repository - MLSMR [45] and 102,634 compounds from the Tuberculosis Antimicrobial Acquisition and Coordinating Facility - TAACF [46], Methods). This demonstrated stable accuracy comparable to our internal test data at 0.74 and 0.75 AUROC for MLSMR and TAACF respectively (Methods). Combined, the MLSMR and TAACF had high within-sample compound similarity (0.76 median maximum pairwise Tanimoto similarity (mmTS), IQR 0.69-0.81). Structural overlap between MLSMR and our internal data was significantly lower (0.56 mmTS, IQR 0.48-0.67, Mann-Whitney U test *p* −*value <* 0.0001, Figure 2d, Supplementary Figure 1e). Training the DMPNN on internal data combined with MLSMR and TAACF data did not improve the AUROC on the test set compared with the model trained only on internal data despite the increase in training data size by 254% (n=325,720 AUROC = 0.73). To further motivate the need for Mtb specific models, we showed that Chemprop models trained on large *E. coli* and *S. aureus* datasets, have substantially lower performance than our models (Supplementary Table 11).

### Prospective eDMPNN hit validation

Mtb are closely related to Streptomyces species and both generally are known to have high levels of intrinsic antibiotic resistance mediated through their hydrophobic waxy cell envelope and other mechanisms [2]. The high level of intrinsic antibiotic resistance has historically challenged growth inhibition prediction for small molecules [47–49]. To prospectively validate the generalizability of the inhibition model to new compounds, we predicted Mtb growth inhibition of 435 diverse molecules (mmTS of 0.41, IQR 0.30-0.51, Methods, Supplementary Methods). Of 435, Fleming predicted 6 to be growth inhibitors and 429 non-inhibitors (Methods). The 435 compounds were then tested on the mc^2^-7000 Mtb strain under conditions similar to the training set growth conditions and drug concentrations. We confirmed 5 of the predicted 6 small molecules were inhibitors of Mtb growth (83.3%), and 382 of the 429 were correctly predicted as not inhibitory (89.0%). Activity of only 48 molecules (11%) from this set was incorrectly predicted (Figure 2e, Supplementary Figure 2e). Not surprisingly, these 48 molecules showed lower similarity to the training data compared to the molecules correctly identified by the eDMPNN (mmTS 0.44 vs. 0.55, Wilcoxon Rank Sum Test, p = 0.03).

### Diffusion extends chemical novelty

We built an equivariant diffusion generative model capable of assembling atomic-level building blocks into *de novo* antibiotic candidates with high structural novelty [34]. The model was trained on 114,933 Mtb drug-like compounds and fragments screening result, conditioning on the experimentally measured level of inhibition (Methods). We then compared the diffusion designed compounds with molecules constructed with SyntheMol, a recently introduced chemical fragment level generative model [33] (Methods). Antibiotic designs generated with diffusion are more unique but also more complex to synthesize compared to designs generated with SyntheMol (6275 designs vs. 710 for diffusion, Wilcoxon Rank Sum Test *p* − *value <* 0.0001, Figure 3a-c). However, synthesis of diffusion designed compounds was predicted to be within the feasible range for the majority of molecules (99% have SAScore *<* 6 and 70% have SCScore *<* 3.5), Methods).

**Figure 3:**
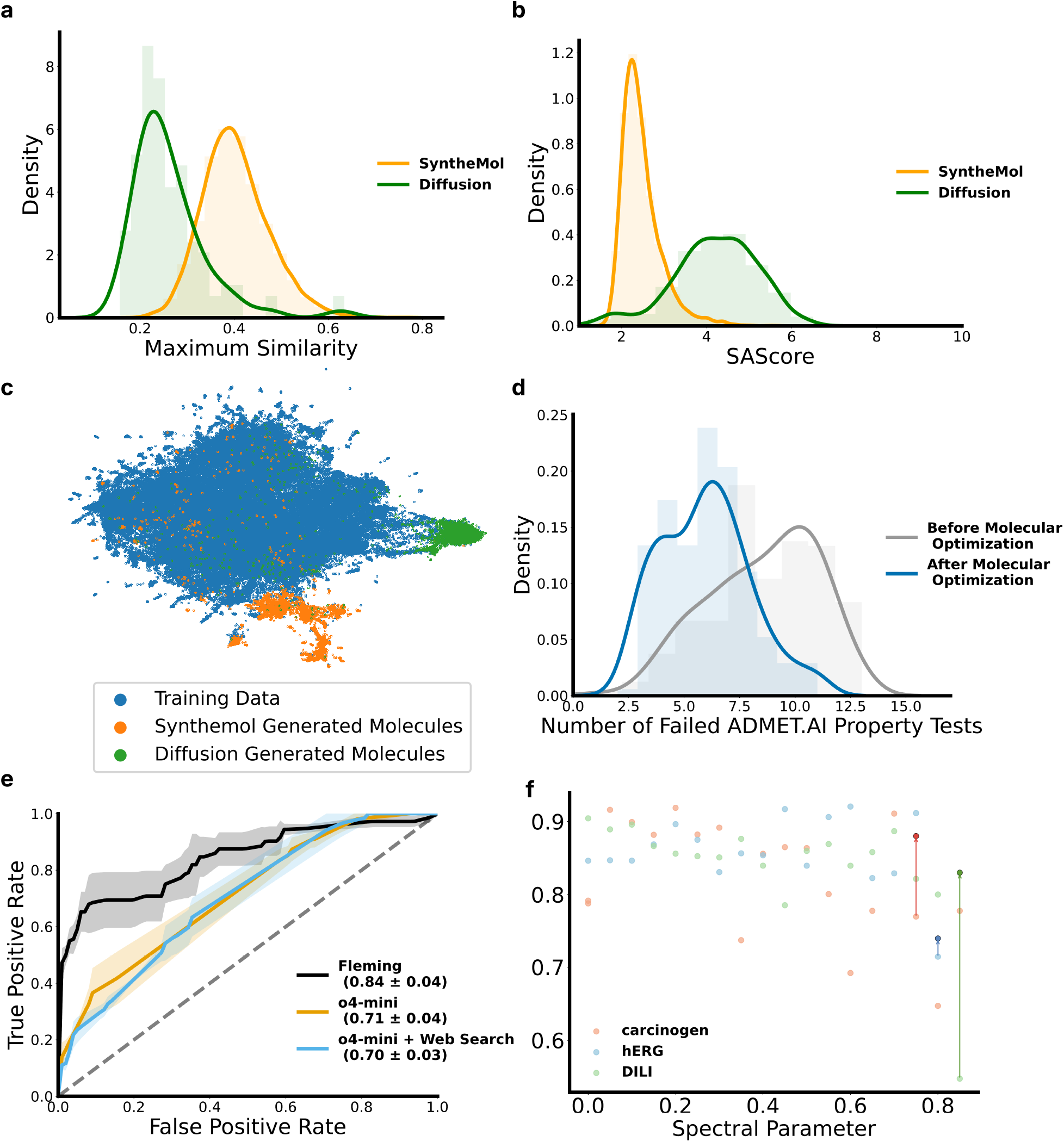
Generation and optimization of novel MTB inhibition hits. **(a)** Distribution of novelty, measured by Tanimoto similarity to the training set, of generated molecules from Synthemol and diffusion model. The lower the similarity, the higher the novelty. **(b)** Distribution of molecular synthesizability, measured by SAScore [61], for Synthemol and the diffusion model. A lower SAScore indicates easier synthesis, with molecules scoring below 6 considered synthesizable [61]. **(c)** A UMAP illustration of structure-based synthesizing method - Synthemol and diffusion models are generating molecules from different sectors of the chemical space. **(d)** Distribution of failed ADMET tasks for molecules before and after optimization by the molecular optimization agent. The fewer failed ADMET tasks, the better. The median number of failed ADMET tasks decreased from 9 (IQR: [7, 11]) to 6 (IQR: [4, 7] after molecular optimization. **(e)** Integrating agents improves the performance of molecular property prediction models. **(f)** Improvement of model performance at high Spectral parameter when integrating Fleming Literature Search.

### Molecular optimization improves ADMET properties of antibiotic designs

The molecular optimization agent is capable of refining initial antibiotic designs using a directed search for molecular neighbors [50] with drug-like properties. While the optimization focuses on ADMET properties, the framework is easily adaptable to other molecular properties provided a property prediction model is available. ADMET profiling is performed using 37 distinct DMPNN models trained on data from the Therapeutics Data Commons (TDC) [51, 52](7 for absorption, 3 for distribution, 5 for metabolism, 3 for excretion and 19 for toxicity) (Supplementary Table 3). We used this agent to optimize 200 designs generated by diffusion or SyntheMol successfully reducing the proportion predicted to fail more than 25% of the ADMET tasks (10/37) from 42.5% to just 5.0% (85 to 10 of 200, Wilcoxon Rank Sum Test *p* − *value <* 0.0001, Figure 3d). Importantly, the optimization process did not alter the predicted inhibition (Supplementary Figure 3d). To further validate the success of the process we used three orthogonal predictors of cytotoxicity (to HepG2, IMR-90 and HSkMC cells in vitro, Methods) that were not used by the molecular optimization agent. Predicted toxicity to HepG2 cells was reduced from 43.1% to 25.4% and to IMR-90 cells from 81.5% to 57.7% after optimization (Fisher *p* − *value* ≤ 0.004). Predicted toxicity to HSkMC cells remained low and did not change significantly, moving from 6.1% to 11.5% (Fisher p-value = 0.19). These results demonstrate the optimization agent’s ability to reliably improve multiple ADMET properties while preserving predicted inhibition of the generative designs.

### Fleming improves generalizability over individual AI models and LLMs

We hypothesized that Fleming is more generalizable to unseen molecules than its individual components *i*.*e*. that it outperforms the sequential molecular AI pipeline without the LLM agent framework or the LLM when used alone (Methods). We tested this by simulating a challenging molecular property prediction task using SPECTRA [43], wherein we constructed training and test sets with sequentially decreasing compound similarity between them. We focused on three ADMET properties – hERG, drug induced liver injury, and carcinogenicity – and found that molecular AI performance predictably decreased as test similarity decreased. While the AUC of the molecular AI model dropped by 30-34% at minimum train-test similarity, Fleming surpassed the AI model with a (4 – 13%) higher AUC at the minimum train-test similarity. Fleming outperformed GPT-4/o4-mini evaluated in isolation (Δ AUC +0.07, Methods, Supplementary Results).

We evaluated Fleming’s ability to distinguish candidate anti-tubercular compounds not yet in clinical use from drug-like molecules with no reported activity against Mtb. Across three independent replicates, Fleming achieved higher accuracy (AUROC = 0.85) than the LLM alone (o4-mini, AUROC = 0.68 and o4-mini with web-search, AUROC = 0.66, Figure 3e), highlighting the advantage of Fleming’s integrative approach.

We also measured impressions of human scientist users of Fleming’s co-pilot functionality, including chemists, structural biologists and biochemists. Scientists were given two cases, an optimization case and a *de novo* design case (Supplementary Table 9); 73% of users rated Fleming highly (*>* 7 on 1-10 likert scale) on meaningful reasoning and validity of the structures (n=11). Additionally, 73% reported they would use the co-pilot again in the future.

### Fleming designed molecules are structurally novel and inhibit Mtb growth *in vitro*

We used Fleming in batch mode to design Mtb inhibitors using diffusion (n = 710 designs) and SyntheMol (n = 6,275). We ranked the top 100 molecules from each method by inhibition, ADMET, novelty and synthesizability, then optimized further for ADMET (Figure 4a). The final candidates were selected using an LLM prompted on whether the design was a plausible antibiotic worthy of experimental testing (Supplementary Notes). Independent human medicinal chemist review of same molecules showed 76% agreement with Fleming’s decision on plausibility.

**Figure 4:**
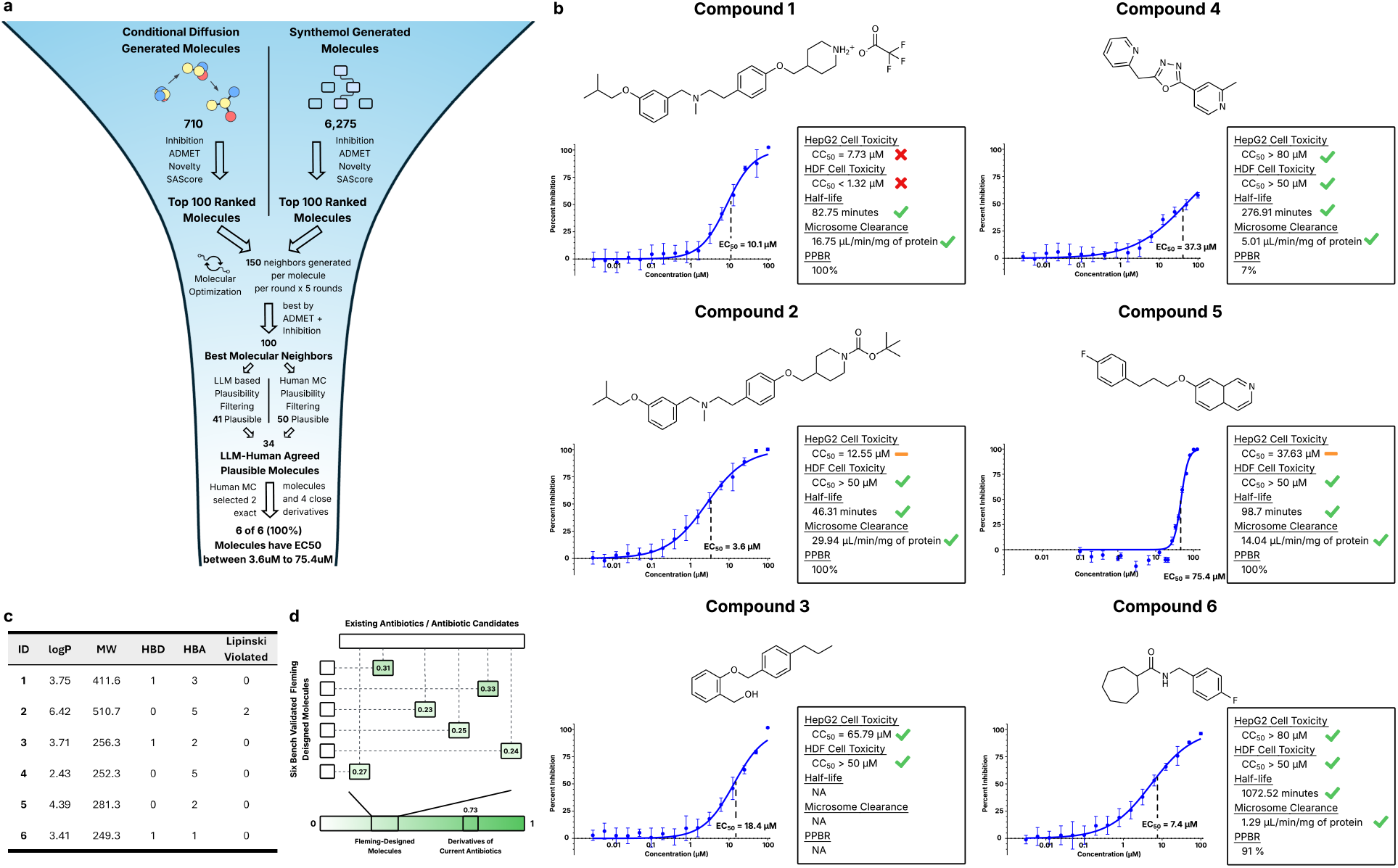
Bench Validation of Fleming Generated Molecules. **(a)** Novel molecules were initially generated using two models: a conditional diffusion generative model and a structure-based generative model (Synthemol). The generated compounds were ranked based on predicted growth inhibition, predicted ADMET properties, molecular novelty relative to the training set, and synthetic accessibility (SAScore). The top 100 molecules from each method were then subjected to five rounds of molecular optimization, guided by predicted inhibition and ADMET profiles, yielding 100 optimized candidates. These were further filtered based on chemical plausibility and lab-specific synthesizability criteria, resulting in six final compounds selected for experimental validation. All six molecules demonstrated significant Mtb growth inhibition in wet-lab assays, with EC50 values ranging from 3.6 *µ*M to 75.4 *µ*M. **(b)** Dose–response curves for the six bench-validated molecules. The x-axis represents compound concentration (*µ*M), and the y-axis shows the percentage inhibition of activation of Mtb growth. The toxicity and pharmacokinetic properties of the candidates, including HepG2 Cell Toxicity, HDF Cell toxicity, human liver microsome half life, human liver microsome clearance are labeled. HEK293Ts data were comparable to the HepG2 cytotoxicity (the HEK293Ts cytotoxicity values could be found in Supplementary Table 10). NA represents the conducted assay is not applicable to the compound. **(c)** Summary table of the six candidate Mtb inhibitors, detailing their inhibitory potency and physicochemical properties. logP here stands for the the logarithm of the octanol-water partition coefficient, MW stands for the molecular weight, HBD and HBA stand for hydrogen bond donors and acceptors, and Lipinski Violated stands for the number of Lipinski rules violated. Five out of six molecules pass all Lipinski’s rules. **(d)** All compounds exhibit low structural similarity to approved antibiotics across microbial species. The structural similarity is calculated by pair-wise maximum Tanimoto Similarity between each of the six candidates and the existing antibiotics/antibiotic candidates.

From the 34 molecules selected by both Fleming and medicinal chemist review, medicinal chemists selected for synthesis based on feasibility. Two were synthesized as designed, and three were modified for synthetic feasibility (retrospectively all three of the modified designs are predicted inhibitory with probability *>* 0.5, Supplementary Figure 4f). For one, the molecule and its N-Boc protected analog were both tested, giving six total molecules (synthesis and structural validation are detailed in the Supplement and Supplementary Figure 4a-e, 5). Remarkably, all six selected molecules inhibited growth of mc^2^-7000 Mtb strain in a dose responsive manner (Methods), with *EC*_50_ ranging from 3.6*µ*M to 75.4*µ*M. This outcome is exceedingly unlikely to occur by chance (probability *<* 10^−10^ to 10^−6^, assuming a random hit rate of 1 to 5%). Five of the six molecules passed all of Lipinski’s Rules [37] (Figure 4c). The six molecules are novel (pairwise Tanimoto similarity *<* 0.33) compared to all molecules in the training set and a database of 11,272 candidate and approved antibiotics (Methods) (Figure 4d,e, Methods).

### Toxicity and Pharmacokinetics Assessment of Fleming-Designed Inhibitors

To further evaluate Fleming’s ability to generate promising Mtb inhibitors for clinical development, we tested the 6 candidates for *in-vitro* toxicity and pharmacokinetics (PK). Five of the 6 candidates demonstrated none to low cytotoxicity to HepG2, HEK293T, and HDF cell lines (Figure 4b, Supplementary Table 10). Since HepG2 was the most sensitive cell line among those tested, we calculated the selectivity indices (SI) using HepG2 cytotoxicity measurements. The N-boc protected analog of the toxic Compound 1 demonstrated selectivity for inhibiting Mtb (Compound 2 SI = 3.5). Of the 4 other molecules, 1 was moderately selective (Compound 4 SI = 2.1), and 2 had high selectivity (Compounds 3 and 6 with SI of 3.6 and 10.8 respectively). Five of the 6 candidates were assayed for human liver microsome clearance and half life (Compound 3 could not be tested due to technical limitations, Methods). All 5 of the assayed compounds demonstrated robust metabolic stability (Figure 4b). Plasma protein binding rates (PPBR) varied from 7-100% comparable to TB drugs in current clinical use (Fig. 4b). Collectively, the moderate to high selectivity, as well as favorable safety and PK profiles observed show that out of the original five generated molecules four performed on the level suitable for hit to lead program candidates. This unprecedented for standard HTS based drug discovery success rate validate Fleming’s ability to accelerate preclinical identification of lead compounds.

## Discussion

We present Fleming, an agentic framework for tuberculosis antibiotic design that integrates a suite of computational drug development tools and AI models, including a state-of-the-art predictor of Mtb growth inhibition. Fleming synthesizes outputs from molecular AI models with relevant knowledge from the scientific literature to produce more generalizable and accurate predictions of molecular properties than its component models alone and other cheminformatic models in the field. Overall, Fleming’s inhibition model demonstrates 17 to 83-fold higher enrichment for *in vitro* Mtb inhibitory activity compared with random drug screening HTS hit rate (1-5%). The LLM agentic module in batch mode effectively reproduces candidate selection by a medicinal chemist. In co-pilot mode, Fleming delivers substantive reasoning and outputs that directly advance antibiotic design. By mitigating common limitations in high-throughput screening including limited chemical novelty, low hit rate, and inadequate optimization of drug-like properties, Fleming will substantially accelerate preclinical lead identification for antibiotic development.

Fleming’s inhibition model, built on an evidential DMPNN, successfully identified Mtb inhibitors across both retrospective external benchmarks and prospective screens. It demonstrated better performance than comparable models, despite a lower AUROC [47, 53, 54]. We attribute this generalizability to the relatively large and diverse training set of 114,933 compounds, the DMPNN architecture, and the evidential framework. In our benchmarking, the diversity of training data was more critical than data scale to model accuracy, as increasing the scale of training data by more than 4-fold by including the less diverse TAACF and MSMLR data did not increase model accuracy. Nevertheless, the eDMPNN has an appreciable false negative rate, and is more likely to miss active molecules that are more distant in chemical space from the training data. Future improvements in molecular representation by including, for example 3-D molecular structure, conformational flexibility and stereochemistry as well as active learning approaches that intentionally screen underexplored regions of the chemical space hold promise for improving molecular property prediction[41, 55, 56].

Flemings conditional diffusion model enabled us to generate a substantially higher number of novel molecules compared to traditional rule-based approaches using molecular substructures. Although computationally expensive and requiring subsequent filtering on chemical validity, diffusion models hold promise to more widely explore the chemical space for molecules of interest.

We synthesized 5 Fleming-generated molecules and 1 derivative with low similarity to training set molecules and found all 6 to be inhibitory of Mtb growth. That all five of the core designs successfully inhibited Mtb in a dose dependent manner is unprecedented and was otherwise expected to occur with lower probability than one in a million. This finding greatly increases enthusiasm for agentic frameworks and generative antibiotic design and their potential for accelerating antibiotic discovery. The generative pipeline’s higher hit rate, along with the strong selectivity and favorable pharmacokinetics of its designed molecules, clearly indicates that ranking and filtering based on predicted ADMET, synthetic accessibility, and literature-grounded plausibility will streamline discovery and increases yield.

Despite these strengths, several area of improvement remain. Incorporating additional constraints on chemical structure during generation could improve yield of synthetically feasible molecules. When inhibition data at multiple compound concentrations become available on diverse chemical sets, it can be used to refine inhibition modeling to enable prediction of relative potency. Lastly, large language models are prone to occasional hallucinations [57], and their substantial computational costs may restrict their adoption in resource-limited research settings. At the moment, open-source models [58–60] have lower performance than leading closed-source models like OpenAI’s o4-mini, but in the near future the field anticipates the release of high-performing open-source models that together with the use of retrieval augmented generation promises to improve utility of predictive tools.

Fleming demonstrates how AI agents will advance drug discovery by unifying molecular generation, property optimization, and literature-based reasoning within a single framework. Although this study centers on TB, the system is easily adapted to other diseases through targeted retraining of component models. Scientists targeting a different disease can simply replace the inhibition training model, extend the generative conditioning labels, or add other property agents (e.g., mechanism-of-action prediction, metabolic stability) without rewriting the orchestration logic. Importantly, we view Fleming as a compliment, not as a replacement for traditional drug discovery pipelines, accelerating molecular design in both well-resourced and under-resourced contexts. In particular, the ability to interact with Fleming through a natural language interface, coupled with open-source components and lightweight inference endpoints, suggests a path toward more inclusive access to drug discovery. As AI systems continue to mature, new agentic frameworks like Fleming will help lower the entry barrier for scientists working on urgent but overlooked global health challenges.

## Data availability

The data used to develop the inhibition prediction agent and molecule generation agent were experimentally tested in-house and are available upon request. The data for training the ADMET agent are publicly accessible through the Therapeutic Data Commons (https://tdcommons.ai/) [51].

## Code availability

The code to run Fleming agent, together with the code to reproduce the results are available on GitHub at https://github.com/farhat-lab/Fleming.

## Acknowledgments

We thank Ryan Hughes, Tabitha Emde, and Jeremy Wood for generating the Mycobacterium tuberculosis growth inhibition data used for model training. A.Z. was supported by T32GM144273 from the National Institute of General Medical Sciences.

## Author contributions

M.F. and Y.E. conceptualized the study. Z.W., Y.E., and A.Z. developed the Fleming agent, trained and evaluated computational models, performed benchmarking, and prepared all figures. D.N., T.D., S.H., I.K., and J.S. supervised and conducted high-throughput screening to generate data for model training. B.A., I.K., J.S., and M.F. provided domain expertise in Mycobacterium tuberculosis antibiotic design. S.H. performed Mtb cell based testing, M.P. synthesized inhibitors, A.F. performed ADME testing, I.K., and J.S. supervised and performed synthesis and experimental validation of Fleming-selected and generatively designed molecules. M.S., A.D.W., and S.R. provided access to and technical support for the PaperQA2 tool, which was tested and integrated as a tool for Fleming. M.Z provided key oversight guiding the DMPNN model and agent development. M.F. supervised the overall project, guided manuscript preparation, and secured funding. Z.W., Y.E., and A.Z. wrote the manuscript with input from all authors. All authors reviewed and approved the final version of the manuscript.

## Methods

### Fleming

Fleming is an AI agent built upon the OpenAI *o4-mini* large language model, that coordinates specialized sub-agents and a suite of tools to provide insights to chemists working on TB drug discovery. Fleming flexibly responds to specific user queries and generates grounded results from specialized component agents, custom-built models, and literature reviews.

There are four primary specialized sub-agents: the growth inhibition agent, ADMET agent, optimization agent, and molecular generation agent. These sub-agents share a subset of general-purpose tools, such as the ability to write and run Python code, search the literature, and use a molecular describer. Each agent also has access to its own collection of tools. The full description of tools and their assignments is presented in Figure 1b.

To respond to the varied demands of medicinal chemists, we also endowed Fleming with the ability to perform large-scale batch processing of molecules, which can be used to analyze either a molecule library the user is interested in, or generate many molecules in one session (See ‘Fleming Batch Mode’).

Agents employ the ReAct paradigm [62], where the agent produces natural language thought processes, tool calls, actions, and responses. All agents generate a high-level plan before beginning any analyses. Then, they are prompted to begin their analysis using their prescribed tools. When a user writes a prompt to Fleming, Fleming decides what tool or sub-agent to invoke. If it is a sub-agent, Fleming autonomously formulates the input instruction.

Each sub-agent concludes its analysis with a detailed summary, which is provided to Fleming. To limit the context window of Fleming, it has access to these summaries in lieu of the full conversational trace [63]. Each agent has full access to its own conversation history. As a result, the manager agent can conduct a continuous conversation with the user, and the specialized component agents can continue from previous analyses if invoked again.

### Specialized Sub-Agents

#### Inhibition Agent

The inhibition agent has access to an Mtb growth inhibition prediction model (See ‘Mtb Growth Inhibition Model’), a rationale extraction module (See Rationale Extraction Module), and the ability to conduct targeted literature reviews on molecules. The agent balances predictions from the Mtb growth inhibition prediction model – which outputs both the inhibition prediction score and the prediction confidence – with other information that it obtains, such as the rationales and findings from the literature review. For instance, if a molecule is predicted with high uncertainty to be inhibitory, but the literature review suggests no inhibitory activity for this class of molecules, the predicted inhibition value will be decreased. The inhibition agent is prompted to return both the raw predicted values from the DMPNN as well as the modified prediction when taking into account all data that it has gathered.

#### ADMET Agent

The ADMET agent is constructed in the same manner as the inhibition agent, but has access to diverse ADMET property prediction AI models (See ‘ADMET Models’) instead of the inhibition prediction models. Since there are many ADMET tasks, the sub-agent has the option to view all the results in a markdown table, or as a binarized summary of number of passed/failed tasks. Like the inhibition agent, it responds to flexible queries from Fleming.

#### Optimization Agent

The optimization agent is tasked with improving the molecular properties of input molecules. In addition to the inhibition and ADMET property prediction models, the agent generates chemical neighbors for given molecules using the SynSpace (v0.3.0) package [50]. In a typical instance, the optimization agent will generate chemical neighbors and evaluate all of them for the property of interest (i.e. hepatocyte toxicity, or TB inhibition). It will select candidates, and conduct literature review on them, before returning the optimized molecules. Crucially, the optimization agent is flexible and can perform this optimization task across user-defined objectives.

#### Molecular Generation Agent

The molecular generation agent employs two generative models: SyntheMol [33] and an equivariant conditional diffusion model [34]. The user can specify either model to generate a potential inhibitory molecule. During the generation process, the agent samples a molecule from the model. Then, it always checks the inhibition prediction using the Inhibition Prediction model. The agent also has access to tools that calculate the novelty, synthesizability, and Brenk/PAINS status.

#### Fleming Batch Mode

Fleming can also run large-scale batch processing. When the user inputs data or requests more than 10 molecules, Fleming runs itself in batch processing mode. Batch mode harnesses the core functionality from the main Fleming, but runs simplified versions of it, and does not permit user interaction.

Fleming batch mode can either take in an input CSV of molecules generate molecules from the generative models upon request. Fleming samples equally from each method and applies *post hoc* filtering to retain molecules with high predicted novelty and synthesizability. For the synthesizability filter, we filter out molecules with an SAScore larger than 6. The novelty filter calculates the Tanimoto Similarities between the output molecules and the inhibition model training data to ensure that generated molecules are not recapitulating the training data. Molecules with similarity greater than 0.3 with existing molecules in the set are removed. Molecules with predicted inhibition score less than 0.7 are also filtered out.

Input or generated molecules are ranked by 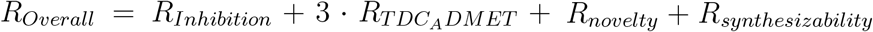 where *R*_*Overall*_ is the ranking used in prioritize the molecules entering the molecular optimization, *R*_*Inhibition*_ is the rank of the predicted inhibition value, 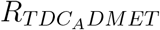 is the rank of the number of passed TDC ADMET tasks, *R*_*novelty*_ is the novelty rank according to maximum similarity against training data, and *R*_*synthesizaibility*_ the SAScore rank.

The user-specified *n* top molecules from each generative model (diffusion or SyntheMol) undergo molecular optimization. In molecular optimization, up to 150 molecular neighbors are generated by Synspace [50]. The top molecular neighbor, defined as the molecule that passes the predicted inhibition threshold (0.7) and fails the fewest TDC ADMET tasks, is assigned to be the starting point of the next round of optimization. The optimization continues for five rounds or until there is no molecular neighbor that fails fewer TDC ADMET tasks than the seed molecule. The *m* best optimized molecules (where *m* ≤ *n*) are taken to the LLM-filtering step. Here, a large language model is prompted to return a binary label of the plausibility of the molecule (Supplementary Notes - Prompts). Those that are deemed plausible are output from the pipeline with their computed molecular properties. These molecules can be further processed and analyzed using core Fleming functionalities.

### Directed Message Passing Neural Network

For the Directed Message Passing Neural Network (DMPNN), the model constructs graph objects based on the molecules’ SMILES strings and generates embeddings using message passing. The molecular embeddings are then concatenated with additional molecular features generated by the RDKit package [64] to predict the Mtb inhibition of the molecules using a multi-layer perceptron. We trained both the classification model and the evidential classification model using the chemprop package (version 1.6.1) [65] with a batch size of 64, depth of 3, and dropout of 0.2. An exponential decay learning rate scheduler is used, with two warm-up epochs, an initial learning rate of 0.0001, a maximum learning rate of 0.001, and a final learning rate of 0.0001.

To identify molecules with the highest likelihood of Mtb inhibition while accounting for noisy training data, we trained DMPNN with evidential classification [44]. Unlike conventional classifiers that output point estimates, evidential models produce uncertainty-aware predictions. By filtering out low-confidence predictions (confidence *<* 0.6), we improved the reliability of the inhibition model’s outputs, resulting in better predictive performance on the retained subset.

### Conditional Equivariant Diffusion Generative Model

We first used RDKit to generate 3D coordinates from 2D SMILES representations, then recalibrated the mapping between atomic distances and bond lengths to maximize coordinate validity. We built on an existing E(3)-equivariant diffusion framework [34], employing an equivariant graph neural network (EGNN) to diffuse over 3D coordinates and atom types. The model was conditioned on each molecule’s measured Mtb inhibition value and trained to generate chemically valid structures. Training used multiple GPUs over three days, with reduced diffusion steps to improve efficiency, a polynomial noise schedule, and normalization to balance geometric, atomic, and property features. During sampling, molecule sizes were drawn from empirical distributions, and only structures passing chemical validity checks were retained.

### Molecular Describer

To enable effective literature queries, we convert the machine-readable SMILES strings to human-readable, natural language molecular descriptions. We first searched two databases for the IUPAC name corresponding to the input SMILES string [66, 67]. If this does not exist in the database, we use Knowledgator Chemical Converters [68] to generate a predicted IUPAC name. We use o4-mini[69] and one chain-of-thought example[70] to generate the description.

### Literature Review Functionality

Fleming, as well as all of the specialized sub-agents, have access to three literature review tools. The agents have access to both a faster literature review (o4-mini with web search) and a deep substructure-based analysis using PaperQA2. Fleming determines which literature review to proceed with based on appropriateness for the task and user preference. For the substructure-based analysis PaperQA2 [71] we used the following procedure: each molecule was processed using the molecular describer tool (See ‘Molecular Describer’) to identify and describe its substructures and functional groups. These descriptors were then distilled into core substructures. Each core substructure was individually input into PaperQA2, which retrieved relevant literature describing the effect of the substructure on the query property. All substructure literature results, along with the molecule’s overall descriptor and smile image, were then provided to o4-mini, which was prompted to assess the molecule’s query property based on literature evidence.

### Mtb Growth Inhibition Dataset

The data we used to train the Mtb inhibition prediction model and the conditional diffusion generative model consists of 113,919 chemically diverse compounds (pairwise Tanimoto similarity *<* 0.65), 1,000 fragments [40], and 14 current in-use TB antibiotics. The compound readouts are from three high-throughput screen runs.

In all cases, mc^2^-7000 strain of Mtb (PMID: 12219086) starter culture was grown in 7H9 media supplemented with OADC, 0.04% Tyloxapol and 25 *µ*g/ml of pantothenate to the OD600 of 1. For the screening, starter culture cells were diluted into testing media to the OD600 of 0.01 and dispensed into 384 well plate.

The first set of 50,000 molecules was tested at a final concentration of 10 *µ*M (compounds were added from 100% DMSO stock, to final 1% DMSO) using M9 minimal media supplemented with 0.5% dextrose, 0.2% sodium acetate (stock solution was pH adjusted to neutral prior to use), 0.01% NaCl, 0.04% Tyloxapol and 25 *µ*g/ml of pantothenic acid. After 5 days of incubation at 37 C, resazurin was added to a final concentration of 0.005 mg/ml to all wells to provide viability readout. Plates were incubated for additional 1-2 days at 37 C, until negative control wells, containing 1% DMSO, were uniformly converted to resorufin pink. Resorufin fluorescence was recorded by OMEGA Polar Star plate reader with excitation wavelength of 540 nm, and emission at 590 nm. Percent inhibition for each compound was calculated using positive (1*µ*M rifampicin) and negative (1% DMSO) controls on each corresponding plate.

The second set was tested at a final concentration of 20 *µ*M (2% DMSO final) in the 7H9 media supplemented with 0.5% dextrose, 0.1% sodium acetate pH 7.0, 0.05% tyloxapol, 25 *µ*g/mL pantothenic acid, and 25 *µ*g/mL kanamycin, using both resazurin and luminescence readouts. For the later, Mtb mc2-7000 strain was transformed with luxCDABE operon containing plasmid enabling constituent expression of both light producing enzyme and the substrate for it (courteously shared with us by Cox lab) [72]. Resazurin conversion to resorufin was recorded as described above, luminescence was measured by OMEGA Clariostar plate reader at 37 C after 5 days of incubation with the compound. Percent inhibition was calculated by normalizing between positive and negative controls included on each plate, as described above.

Finally, a set of 100,000 molecules, partially overlapping with the first set, was tested under the same growth conditions as set two, using luciferase expressing mc^2^-7000 strain, and recording luminescence as a viability readout.

The 1000 compound fragment collection (Maybridge Ro3 library, MW *<* 300 g/mol) was tested against the Mtb H37Rv strain in complete Middlebrook 7H9 broth, with optical density (OD) at 600 nm as readout as described [40]. Due to the anticipated weak inhibitory effects of fragments, 1 mM was used in a single-point screen [40].

We binarized inhibition using an internally calibrated threshold. Molecules with normalized inhibition at or more than 40% of the corresponding plate controls are labeled as inhibitors, while other molecules are labeled as non-inhibitors. Molecules with replicates are labeled as inhibitory if any replicate that exceeds 40% inhibition. In the fragment dataset, fragments were classified as inhibitory if they reduced Mtb growth by at least 70% relative to the drug-free control at a screening concentration of 1 *µ* M.. All current TB antibiotics were labeled as inhibitors when added to the training data.

### Benchmarks and Evaluations

#### Mtb Growth Inhibition Model Validation

For each model architecture, we trained five separate models with 80%/10%/10% scaffold-balanced [73] splits for training, validation, and test sets, grouping molecules by scaffold created by the Chemprop package [65] to prevent data leakage and ensure robust evaluation. The mean Area Under the Receiver Operating Characteristic Curve (AUROC) and Area Under Precision Recall Curve (AUPRC) are reported to evaluate the performance of the models. In the plots, error bars represent the standard deviations of the model performances of the five models. For the final model deployment, we retrained a set of three replicates on the entire dataset (90% train, 10% validation). We also measured the area under the performance-versus-spectral-parameter curve (AUSPC) using the spectral framework for model evaluation (SPECTRA) [43] that generates dataset splits based on a spectral parameter ranging from 0 to 1. Higher spectral parameters (closer to 1) produce splits with lower train-test similarity, while lower spectral parameters (closer to 0) result in splits with higher train-test similarity. Tanimoto similarity was used as the metric to evaluate similarity between the train and test set. AUSPC quantifies the model’s generalizability across varying spectral parameter values. We trained and evaluated each model on splits generated using spectral parameters ranging from 0.0 to 0.75. The maximum spectral parameter was determined by maintaining the training dataset size of at least 1,000 molecules, and the maximum spectral parameter indicates the inclusion of all molecules in the original training dataset.

AUROC, AUPRC, and AUSPC were computed for all benchmarked models on the entire test set, with the exception of the evidential DMPNN. For the evidential DMPNN, performance metrics were calculated only on test set predictions with a confidence score exceeding 0.6.

#### MLSMR and TAACF Evaluation

We obtained the MLSMR downloaded the MLSMR dataset from Pubchem [74] (assay AIDs: 1332, 1626). The TAACF-CB2 dataset [75] and TAACF-Kinase datasets were downloaded from Pubchem (assay AID: 1949 and 2842, respectively). The datasets were concatenated to create a single MLSMR and single TAACF dataset. We dropped all duplicate molecules from each of these datasets.

We calculated all pairwise Tanimoto similarities within a dataset of interest. The value reported for a given molecule is the maximum Tanimoto similarity within its dataset. For model evaluation, we constructed random train-test splits for the the MLSMR and TAACF datasets. We evaluated our trained growth inhibition model on these data (Mtb Growth Inhibition DMPNN). We then trained a Chemprop model on our train set supplemented with the MLSMR and TAACF train sets. We used the same hyperparameters as when training our model (Mtb Growth Inhibition DMPNN). We evaluated performance on our test set.

To visualize the difference between these datasets, we computed the Morgan Fingerprints (radius=2, 2048 bits) for all molecules in our dataset, the MLSMR, and the TAACF sets. We performed dimensionality using Uniform Manifold Approximation and Projection (UMAP, scikitlearn version 1.5, [76], n neighbors=100, min dist=0.1, metric=Jaccard).

#### Evaluation of Diffusion Model and Synthemol

We evaluated the synthesizability and novelty of molecules generated by the diffusion model (710 molecules) and SyntheMol (6,275 molecules).

Synthesizability was evaluated using the Synthetic Accessibility Score (SAScore), which estimates the ease of chemical synthesis. To assess novelty, we computed the mean maximum pairwise Tanimoto similarity (MMPT) between each generated molecule and all molecules in the training set. Lower MMPT values indicate greater structural dissimilarity and, hence, higher novelty.

#### Molecular Optimization Evaluation

We selected 200 molecules and passed them through the molecular optimization agent with the objective of minimizing the total number of failed ADMET prediction tasks from the Therapeutics Data Commons (TDC). Of these, 130 molecules were successfully optimized. To evaluate optimization performance, we compared the mean number of failed TDC ADMET tasks before and after optimization. We also evaluated the three cytotoxicity properties of the molecules that had been successfully optimized before and after molecular optimization using external models (See Method - ADMET Models). The predicted values are binarized using 0.7 as the cut-off.

#### Novelty of Generated Molecules

To evaluate the novelty of the generated molecules. We calculate the similarity between the designed molecules and existing antibiotics and antibiotic candidates. The existing antibiotics are collected by searching ChEMBL and collected molecules under ATC codes: “Antiinfectives for Systemic Use (J01)”, “Antivirals for Systemic Use (J05)”, “Antimycobacterials (J04)”, “Antibiotics and Chemotherapeutics for Dermatological Use (D06)”, “Antimycotics for Systemic Use (J02)”, “Gynecological Antiinfectives and Antiseptics G01)”, “Ophthalmologicals (S01)” and “Otologicals (SO2)”. Together, these comprise 11,272 unique molecules. The detailed list of antibiotic is included in Supplementary Table 4 We evaluated the similarity between the designed molecules and this set using the Tanimoto similarity. We reported the maximum Tanimoto similarity between the molecule of interest and all of the molecules in this set. For reference, we calculated the pairwise Tanimoto similarity among the antibiotic class Penicillin. We additionally collected the small molecules with publicly available SMILES in the Preclinical and Phase 1 studies [77], and calculated their similarity to the ChEMBL-derived molecules.

#### Fleming Improves Generalizability of ADMET Models

We first applied SPECTRA to the DILI, Carcinogen, and hERG datasets from the Therapeutics Data Commons (TDC)[51]. For each dataset, we selected splits corresponding to the highest spectral parameter where the subset size exceeded ten molecules, ensuring sufficient data for model training. We then trained and evaluated Chemprop models on each SPECTRA split. We focused on the two most challenging conditions, those with the highest spectral parameters, and evaluated Fleming in a zero-shot setting on all molecules in the test sets. In addition, we ran a variant of Fleming in which all auxiliary tools were removed except the molecular describer. Importantly, the original Fleming had access to the Chemprop model trained specifically on the corresponding split, avoiding leakage that would arise from using a general Chemprop model trained across all data. Predictions from each model were used to compute the AUC for comparative analysis between the chemprop model, Fleming, and modified Fleming.

#### Fleming Distinguishes TB Drugs from Control Molecules

We created a benchmark for Fleming that tests its ability to reason over multiple molecular properties. We selected all small molecules in preclinical, Phase 1, and Phase 2 studies that had public structures [77], reasoning that these molecules should have TB inhibitory properties and a favorable ADMET profile. We excluded drugs that had a Tanimoto Similarity greater than 0.65 to other drugs in the set.

To prepare a list of drug-like molecules without anti-TB activity, We collected approved small-molecule drugs listed in Chembl [78]. We filtered this down by removing molecules with fewer than 6 heavy atoms and drugs that have been discontinued. To reduce the chance of any of these control compounds being active against TB, we only included drugs from the following classes: Glucocorticoid, Antihypertensive, Progestin, Estrogen, Bronchodilator, Vasodilator (coronary), Analgesic (narcotic), Diuretic, Antihypertensive; Diuretic, Adrenergic (vasoconstrictor), Antihyperlipidemic, Antitussive, Vitamin (enzyme co-factor), Vitamin (prothrombogenic), Oxytocic, Adrenocortical Steroid (salt-regulating). We then subset these molecules to those that are known *invitro* noninhibitors[79]. Finally, we excluded molecules that had a Tanimoto Similarity greater than 0.65 when compared to other molecules in the set.

Our benchmark thus is comprised of anti-TB candidates labeled as inhibitors, and the collected non-TB drugs labeled as noninhibitors. All molecules selected are included in the Supplementary Table 6. We then prompted Fleming, o4-mini, and o4-mini equipped with web-search to distinguish between the control molecules and the TB candidates (Figure 3e). Prompts are available in the Supplementary Notes. We ran three replicates for each model.

### *In-vitro* Library Screening Validation

To prospectively validate our inhibition prediction model, we used a diverse chemical set from another collaborative project: commercially available compounds comprising Enamine REAL and HTS collections and MCule collection were virtually screened by CHARM Therapeutic by cofolding with two Mtb targets: Pks13 TE domain and PknB. These compounds were used to test prediction by Fleming and wet lab evaluation against Mtb (Supplementary Figure 2a,b). For consistency, we used the same experimental conditions as in the second batch of training data. The mc2-7000 strain of Mtb (PMID: 12219086) starter culture was grown in 7H9 media supplemented with OADC, 0.04% Tyloxapol and 25 *µ* g/ml of pantothenate to a OD600 of 1. Starter culture cells were diluted into testing media to a OD600 of 0.01 and dispensed into 384-well plates. Candidates are tested at a final concentration of 100 *µ* M in the 7H9 media supplemented with 0.5% dextrose, 0.1% sodium acetate pH 7.0, 0.05% tyloxapol, 25 *µ* g/mL pantothenic acid, and 25 *µ* g/mL kanamycin, using resazurin readouts.

We passed the molecules from the library through eDMPNN and categorized the molecules as hits if the predicted inhibition *>* 0.7 and prediction confidence *>* 0.6. The distribution of the prediction is provided in the Supplement (Supplementary Figure 2d).

We also benchmarked the performance of the selected model architectures on the screened library. eDMPNN still shows the highest AUROC compared to other architectures (Supplementary Figure 2c).

### *In-vitro* Validation of Fleming-designed molecules

To evaluate Fleming’s performance in designing novel Mtb antibiotics, we tested the molecules generated from Fleming *in-vitro*. Since Fleming was tasked with exploring a large set of molecules, batch mode was invoked (see Methods Batch Mode). We generated 6,985 predicted inhibitors, 710 from the conditional diffusion generative model, and 6,275 from SyntheMol [33]. Using the Batch Mode protocols, we filtered each set down to the top 100 molecules. We then ran the batch mode molecular optimization pipeline and selected the top 100 best molecular neighbors. Finally, we conducted an LLM-based plausibility filtering step ([69]). We prompted a GPT-4o instance to assess the plausibility of the molecules as antibiotics (Supplementary Method). We also independently asked medicinal chemists to rate the molecules’ plausibility. The molecules that were rated as plausible by both the LLM and the humans were retained. Among these molecules, human medicinal chemists selected six molecules to synthesize and validate experimentally based upon synthesizability and feasibility within their laboratory conditions. In the end, the chemists synthesized two molecules as-is, and four close to initial generative designs analogs (See Supplementary Figure 4f).

#### Experimental Setting for Candidate Molecule Validation

The experimental settings for testing the Fleming-designed molecules are the same as that of the *Invitro* library screening validation. Candidates are tested in a dose response with the concentration of 100 *µ*M in the 7H9 media supplemented with 0.5% dextrose, 0.1% sodium acetate pH 7.0, 0.05% tyloxapol, 25 *µ*g/mL pantothenic acid, and 25 *µ*g/mL kanamycin, using resazurin readouts.

