## Supplementary Materials for "Fleming: An AI Agent for Antibiotic Design for Mycobacterium tuberculosis"

This PDF file includes:

Supplementary Notes
Supplementary Figures
Supplementary Tables

### S1 Supplementary Notes

#### S1.1 Supplementary Results

##### S1.1.1 Fleming Improves Generalizability of ADMET Models

Running SPECTRA on the DILI, Carcinogen, and hERG datasets revealed a degradation in performance as the spectral parameter increased, consistent with prior observations of reduced generalizability under high spectral separation [1]. We focused on spectral parameters of 0.8 and 0.85, which exhibited the largest AUC declines relative to the baseline at parameter 0. Fleming substantially improved performance in these challenging regimes—for example, increasing AUC from 0.80 to 0.90 at SP = 0.8 and from 0.55 to 0.83 at SP = 0.85 for DILI, from 0.71 to 0.74 at SP=0.8, from 0.93 to 0.97 at SP=0.85 for hERG and from 0.65 to 0.73 at SP = 0.8 for Carcinogen, while showing a decrease (0.78 to 0.69) at SP = 0.85. The full Fleming system outperformed the stripped variant containing only the molecular describer in the DILI and Carcinogen datasets, it did not outperform the stripped variant in hERG. We hypothesize this reflects dataset-specific memorization effects, as the model appeared to recognize previously encountered molecules based on their SMILES representations. These results highlight that Fleming provides complementary reasoning capabilities beyond conventional ML models or language models alone, particularly in high-divergence regions where standard predictors exhibit poor generalization due to minimal structural overlap between train and test compounds.

#### S1.2 Supplementary Methods

##### S1.2.1 Benchmarked Models

**Equivariant Graph Neural Network** The Equivariant Graph Neural Network (EGNN) [2] is a model architecture equivariant to translation, rotation, reflection, and permutation, leveraging the 3D positional data of each atom in the molecule as input. We used a batch size of 64, a hidden layer dimension of 64, 3 layers, gradient clipping at 0.2, and no dropout. Training employed an exponential decay learning rate scheduler with 4000 warm-up steps. We used a grid search to determine optimal hyperparameters.

**Transformer-M** Transformer-M [3] is a state-of-the-art model that uses both 2D and 3D information of a molecule as model input. It incorporates a self-supervised 3D Positional Denoising objective [4] in the 3D channel as a co-objective to enhance model performance. We used a grid search to determine optimal hyperparameters. We used a batch size of 256, a noise scale of 0.2, and a channel weight of [1,0.01]. We also used an exponential decay learning rate scheduler with an initial learning rate of 1e-5.

**Feed Forward Neural Network** We used a Feedforward Neural Network (FFNN) with RDKit Fingerprints to test the effectiveness of RDKit fingerprints alone in predicting the Mtb inhibition. We used a grid search to determine optimal hyperparameters. We used a 10-layer FFNN with a batch size of 64, a hidden layer dimension of 256, a gradient clip of 0.2, and no dropout. An exponential decay learning rate scheduler is used with 4000 warm-up steps.

**Random Forest** We used a random forest model from scikit-learn (version 1.5) [5] with 100 decision trees. The input to the random forest model is the RDKit fingerprints, which is the same as the input of the FFNN.

##### S1.2.2 Rationale Extration Module

For a given molecule, the Rationale Module finds the molecular substructure that drives the model's prediction and assesses whether this rationale is likely to be the core inhibitory element of an antibiotic. To do this, we employ a Monte Carlo Tree Search to identify subgraphs of the molecules that are between $S_{min}$ and $S_{max}$ atoms with predicted inhibition above $\delta$ [6]. This method has been used previously to identify rationales of antibiotic inhibition[7] and is implemented in Chemprop (version 1.6.1) [8]. We used parameters: $\delta = 0.5, S_{min} = 8, S_{max} = 14, C_{PUCT} = 10$, rollouts=20. We use the rationale with the highest inhibition scores as the final rationale for the molecule.

##### S1.2.3 ADMET Models

We selected 37 ADMET-AI models that were trained on datasets from the Therapeutic Data Commons (TDC) [9] to predict ADMET properties related to Mtb Antibiotic Discovery. Checkpoints were used from the ADMET-AI suite of models [10].

Additionally, we trained DMPNNs to predict cytotoxicity in human liver carcinoma cells (HepG2), lung fibroblast cells (IMR-90), and primary skeletal muscle cells (HSkMC). We used screening data from 39,312 compounds reported in a previous publication [7]. Our models (trained with Dirichlet Loss, evidential regularization = 0.2, dropout = 0, 200 epochs, learning rate 0.0001) had excellent performance on the test sets ($AUROC_{HepG2} = 0.84, AUROC_{HSkMC} = 0.92, AUROC_{IMR-90} = 0.82$). We also used prebuilt Uni-Mol models [11, 12] trained on MoleculeNet [13] datasets to predict side effects and solubility.

The predicted properties are then binarized as pass or fail by comparing them to the predictions for currently used antibiotics generated by the same model. Screening molecules with predicted property values in the top 90% of values observed for anti-TB antibiotics are classified as pass; otherwise, they are classified as fail. Detailed TDC ADMET properties, corresponding thresholds, and the antibiotics used in setting the threshold are provided in the supplement (Supplementary Table 2,3).

To provide a more comprehensive view of the ADMET profiles, we also collected a Blood Brain Barrier Prediction Model, a side effect prediction model, and solubility prediction model[11] based on a fine-tuned Uni-Mol [14] model and incorporated them as tools for the ADMET agent.

##### S1.2.4 Cytotoxicity Prediction Models

We collected cytotoxicity labels for 39,312 molecules on three human cell lines (HepG2,IMR-90, and HSKMC)[7]. We trained Chemprop models on each, using a random data split (80% train, 10% validation, 10% test). We used the same hyperparameters as our model (DMPNN Growth Inhibition Model). The performance on the test splits were as follows: HepG2 AUROC = 0.84, HSKMC AUROC = 0.92, IMR-90 AUROC = 0.82.

##### S1.2.5 SyntheMol

We used SyntheMol [15], which is a Monte Carlo Tree Search-based package to generate synthesizable molecules with Mtb inhibition. Within SyntheMol, scores related to the selected property of the 132,000 building blocks are predicted by our inhibition model. Molecules are then generated from the building blocks through 13 valid chemical reactions based on inhibition scores. The model then iteratively builds molecules that are likely Mtb inhibitors. The default set of hyperparameters [15] is used for generating Mtb growth inhibitory molecules in the molecule generation agent.

###### S1.2.6 Chemical Space Visualization

For chemical space visualization in this study, we used UMAP [16] to reduce the Morgan Fingerprints (nBits=2048) [17] with Jaccard distance into two dimensions. Matplotlib v2.9.0 [18] and Plotly v.5.23.0 [19] are used in visualizing the chemical space.

###### S1.2.7 Molecule Similarity Calculation

Tanimoto similarity was used to assess molecular similarity. All calculations were performed using Morgan fingerprints with a radius of 2 and a vector size of 2048 bits.

###### S1.2.8 Novelty Calculation

We calculated novelty by finding the maximum pairwise Tanimoto similarity of the given molecule to a combined set of the Chembl antibiotics and our training dataset. In batch processing mode, the novelty is only calculated with respect to the training set.

###### S1.2.9 Synthesizability Calculation

We evaluated molecular synthesizability using two complementary metrics: the Synthetic Accessibility Score (SAScore) [20] and the Synthetic Complexity Score (SCScore) [21]. SAScore values were computed using the RDKit cheminformatics toolkit [22], with molecules considered synthesizable when their SAScore was below 6, consistent with the original definition [20]. SCScore, a neural network–based metric estimating synthetic feasibility, was also calculated, and molecules with SCScore values below 3.5 were classified as synthesizable following prior studies [21, 23].

###### S1.2.10 Morgan Fingerprint

To obtain a numerical representation of molecules, we used Morgan fingerprints, a widely used circular fingerprint [24]. Unless otherwise specified, we computed fingerprints with radius 2 and a 2048-bit vector size. Fingerprints are calculated using RDKit [22].

###### S1.2.11 PAINS Criteria

Pan-Assay Inteference Compouds (PAINS) are substructures that are often found to be false positives in high-throughput screens [25]. We used the RDKit PAINS filters [22] to assess whether molecules possess these substructures. If they do not, we deem them as having met PAINS criteria.

###### S1.2.12 RDKit Documentation Search

To enable more effective use of RDKit functions, we built a retrieval-augmented generation pipeline over the RDKit Documentation. We downloaded the RDKit documentation (2019.09.01), and chunked it by subsection. We created and stored vector embeddinsg of each of these sections using OpenAI's text-ada-002 model. During retrieval, we find and return the four most similar sections to a given input query.

###### S1.2.13 Synthesis of Fleming Designed Molecules

**Compound 1: N-(3-isobutoxybenzyl)-N-methyl-2-(4-(piperidin-4-ylmethoxy)phenyl)ethan-1-amine** To a stirred solution of the 4-(2-((3-isobutoxybenzyl)(methyl)amino)ethyl)phenol (Compound 2) (37 mg, 0.07 mmol) in dry dichloromethane (4 mL) was added TFA (0.1 mL, 1.5 equiv) at 0 °C. The reaction mixture was allowed to warm to room temperature and starred for 2 hours. After completion of the reaction, the reaction mixture was concentrated and crude reaction mixture was dissolved in ethyl acetate, neutralized with aqueous solution of sodium bicarbonate, washed with water (5 mL), and extracted with ethyl acetate (3 × 10 mL). The combined organic layers was washed with brine, dried over anhydrous sodium sulfate, filtered, concentrated and reaction mixture was purified by column chromatography (2% methanol in dichloromethane) to provide the desired product in 60% isolated yield (Figure S5).

**Compound 2: *tert*-butyl 4-((4-(2-((3-isobutoxybenzyl)(methyl)amino)ethyl)phenoxy)methyl)-piperidine-1-carboxylate** To a stirred solution of the 4-(2-((3-isobutoxybenzyl)(methyl)amino)-ethyl)phenol (1) (97 mg, 0.311 mmol) in dry acetone (6 mL) was added potassium carbonate (86 mg, 0.622 mmol) under an inert atmosphere. A solution of tert-butyl 4-(bromomethyl)piperidine-1-carboxylate (2) (95 mg, 0.34 mmol) in dry acetone (2 mL) was added to it. The reaction mixture was starred at 70 °C for 5 hours. After completion of the reaction, the reaction mixture was cooled to room temperature, potassium carbonate was filtered and concentrated. Crude reaction mixture was dissolved in ethyl acetate, washed with water (5 mL), and extracted with ethyl acetate (3×5 mL). The combined organic layers was washed with brine, dried over anhydrous dried over sodium sulfate, filtered, and concentrated. The crude product was purified by column chromatography (1% methanol in dichloromethane) to provide the desired product in 53% isolated yield (Figure S5).

**Compound 3: *tert*-butyl 4-(((2-((4-propylbenzyl)oxy)benzyl)oxy)methyl)piperidine-1-carboxylate** To a stirred solution of 2-(hydroxymethyl)phenol (3) (80 mg, 0.64 mmol) in acetonitrile (5 mL) under an inert atmosphere $K_2CO_3$ (180 mg, 1.30 mmol) followed by 1-(bromomethyl)-4-propylbenzene (4) (151 mg, 0.70 mmol) were added at room temperature and stirred for 8 hours at 60 °C. Upon completion, the reaction mixture was cooled with room temperature, filtered and concentrated. Crude reaction mixture was dissolved in ethyl acetate, diluted with water (5 mL), and extracted with ethyl acetate (3×8 mL). The combined organic layers was washed with brine, dried over anhydrous $Na_2SO_4$, filtered, and concentrated. The crude product was purified by column chromatography (4:1-hexanes/EtOAc) to obtain the desired product in 65% isolated yield (Figure S5).

**Compound 4: 2-(2-methylpyridin-4-yl)-5-(pyridin-3-ylmethyl)-1,3,4-oxadiazole** To a stirred solution of the 2-methylisonicotinic acid (5) (46 mg, 0.33 mmol) and 2-(pyridin-2-yl)acetohydrazide (6) (58 mg, 0.38 mmol) in anhydrous DCM (5 mL) was added triethylamine (0.16 mL, 1.00 mmol) at 0 °C. Propylphosphonic anhydride ($T_3P$, 50% w/w solution in EtOAc, 0.33 mL, 2.0 equiv) was added dropwise. The reaction mixture was stirred at 0 °C for 10 min and then stirred at room temperature for 1 hour. Upon completion, water was added to it and extracted with DCM. The combined organic layers was washed with brine, dried over $Na_2SO_4$, filtered, and concentrated. The crude residue was purified by silica gel column chromatography (2% methanol in dichloromethane) to obtain the desired product (7) in 56% yield. In next step, crude reaction mixture of 2-methyl-N′-(2-(pyridin-3-yl)acetyl)isonicotinohydrazide (7) was dissolve in THF (5 mL), $Et_3N$ (0.15 mL, 1.10 mmol), followed by p-toluene sulfonyl chloride (70 mg, 0.37 mmol) was added to the solution at 0 °C, and stirred for 6 hours at room temperature. Reaction was quenched with water, extracted with DCM, concentrated, and the crude product was purified by silica gel column chromatography (2% methanol in dichloromethane) to provide the desired product in 57% yield (Figure S5).

**Compound 5: 7-(3-(4-fluorophenyl)propoxy)isoquinoline** To a stirred solution of the 3-(4-fluorophenyl)propan-1-ol (8) (0.50 mg, 0.32 mmol) equiv) and Isoquinolin-7-ol (9) (0.043 g, 0.32

mmpl) in dry THF (5 mL) triphenylphosphine (0.112 g, 0.48 mmol), followed by diethyl azodicarboxylate (DEAD), (0.25 mL, 0.49 mmol 40% solution in toluene) was added under an inert atmosphere. The reaction mixture was allowed to warm to room temperature and stirred for 5 hours. Reaction progress was monitored by TLC. Upon completion, the reaction mixture was concentrated and the crude reaction mixture was dissolved in ethyl acetate, diluted with water (5 mL), and extracted with ethyl acetate (3 × 10 mL). The combined organic layers was washed with brine, dried over anhydrous sodium sulfate, filtered, and concentrated under reduced pressure. The crude product was purified by column chromatography (2% methanol in dichloromethane) to obtain the desired product in 56% isolated yield (Figure S5).

**Compound 6: *N*-(4-fluorobenzyl)cycloheptanecarboxamide** To a stirred solution of the cycloheptanecarboxylic acid (10) (47 mg, 0.33 mmol) and 4-Fluorobenzylamine (11) (46 mg, 0.36 mmol) in anhydrous dichloromethane (4 mL) was added triethylamine (0.3 mL, 4.0 equiv) under an inert atmosphere at 0 °C. Propylphosphonic anhydride ($T_3P$, 50% w/w solution in EtOAc, 0.34 mL) was added dropwise over 5 min. The reaction mixture was stirred at 0°C for 10 min and then allowed to warm to room temperature and stirred for 1 hour. Upon completion (monitored by TLC), the reaction was quenched with water. The aqueous layer was extracted with DCM and water. The combined organic layers was washed with brine, dried over $Na_2SO_4$, filtered, and concentrated under reduced pressure. The crude residue was purified by silica gel column chromatography (4:1-hexanes/EtOAc) to obtain the desired product in 29% yield (Figure S5).

##### S1.2.14 Synthesized Compound Validation using $^1$H NMR

**Compound 1** $^1$H NMR (400 MHz, CDCl3): $\delta$ 7.30 (t, J = 7.85 Hz, 1H), 7.10-7.02 (m, 3H), 6.98-6.94 (m, 1H), 6.91 (d, J = 7.73 Hz, 1H), 6.83 – 6.77 (m, 2H), 4.39 – 4.25 (m, 1H), 4.13 – 3.00. (m, 2H), 3.79 (d, J = 5.71 Hz, 2H), 3 .71 (d, J = 6.55 Hz, 2H), 3.51– 3.32 (m, 3H), 3.10 – 2.85 (m, 5H), 2.75 (s, 3H), 2.12 – 1.96 (m, 4H), 1.78 –1.64 (m, 2H), 1.01 (d, J = 6.75 Hz, 6H); $^{13}$C NMR (100 MHz, CDCl3) $\delta$ 160.10, 157.88, 130.27, 129.80, 128.13, 122.64, 121.02, 116.97, 116.10, 114.97. 74.52, 71.20, 59.94, 56.82, 43.71, 39.24, 34.18, 29.63, 28.24, 25.53, 19.09); MS (LCMS): m/z 411.17 [M+H]+.

**Compound 2** $^1$H NMR (400 MHz, CDCl3): $\delta$ 7.18 (t, J = 7.73 Hz, 1H), 7.10-7.05 (m, 2H), 6.89-6.82 (m, 2H, 6.81 – 6.75 (m, 3H), 4.22 – 4.09 (m, 2H), 3.76 (d, J = 6.52 Hz, 2H), 3 .70 (d, J = 6.53 Hz, 2H), 3.53 (s, 2H), 2.80 – 2.68 (m, 4H), 2.66 – 2.59 (m, 2H), 2.29 (s, 3H), 2.13– 2.00 (m, 1H), 1.99 – 1.87 (m, 1H), 1.84 – 1.76 (m, 2H), 1.46 (s, 9H), 1.32– 1.88 (m, 2H), 1.02 (d, J = 6.70 Hz, 6H); $^{13}$C NMR (100 MHz, CDCl3) $\delta$ 159.40, 157.34, 154.88, 140.41, 132.56, 129.62, 129.07, 121.14, 114.98, 114.38, 113.28,. 79.35, 74.36, 72.40, 62.13, 59.29, 42.20, 36.26, 32.88, 28.93, 28.48, 28.35, 19.30; MS (LCMS): m/z 511.14 [M+H]+.

**Compound 3** $^1$ NMR (400 MHz, CDCl3): $^1$ NMR (400 MHz, CDCl3): $\delta$ 8.65 (d, J = 5.16 Hz, 1H), 8.58 (d, J = 5.16 Hz, 1H), 7.76 (s, 1H), 7.72 – 7.65 (m, 2H), 7.36 (d, J = 7.53 Hz, 1H), 7.25 – 7.21 (m, 1H), 4.51 (s, 2H), 2.64 (s, 3H); $^{13}$C NMR (100 MHz, CDCl3) $\delta$ 163.77, 159.87, 150.15, 149.89, 137.12, 131.25, 123.24, 122.68, 119.89, 117.50, 34.65, 24.46; MS (LCMS): m/z 252.85 [M+H]+.

**Compound 4** $^{1}$ NMR (400 MHz, CDCl3): $^{1}$ NMR (400 MHz, CDCl3): $\delta$ 8.65 (d, J = 5.16 Hz, 1H), 8.58 (d, J = 5.16 Hz, 1H), 7.76 (s, 1H), 7.72 – 7.65 (m, 2H), 7.36 (d, J = 7.53 Hz, 1H), 7.25 – 7.21 (m, 1H), 4.51 (s, 2H), 2.64 (s, 3H); $^{13}$C NMR (100 MHz, CDCl3) $\delta$ 163.77, 159.87, 150.15, 149.89, 137.12, 131.25, 123.24, 122.68, 119.89, 117.50, 34.65, 24.46; MS (LCMS): m/z 252.85 [M+H]+.

**Compound 5** $^{1}$ NMR (400 MHz, CDCl3): $\delta$ 9.13 (bs,1H), 8.42 (bs, 1H), 7.72 (d, J = 8.63 Hz, 1H), 7.59 – 7.53 (m, 1H), 7.38 – 7.32 (m, 1H), 7.21 – 7.13 (m, 3H), 7.01 – 6.93 (m, 2H), 4.08 (t, J = 6.33 Hz, 2H), 2.83 (t, J = 7.47 Hz, 2H), 2.21 – 2.10 (m, 2H). $^{13}$C NMR (100 MHz, CDCl3) $\delta$ 162.61, 160.19, 157.75, 151.04, 141.24, 136.88, 136.85, 131.39, 129.87, 129.79, 128.06, 123.72, 120.24, 115.32, 115.11, 105.55, 66.95, 31.35, 30.76; LCMS: m/z 281.89 [M+H]+.

**Compound 6** $^{1}$ NMR (400 MHz, CDCl3): $\delta$ 7.27-7.19 (m, 2H), 7.03-6.97 (m, 2H), 5.71 s, 1H), 4.38 (d, J = 5.82 Hz, 2H), 2.29-2.19 (m, 1H), 1.95-1.84 (m, 2H), 1.82 –1.62 m, 4H), 1.61-1.38 (m, 6H) ; $^{13}$C NMR (100 MHz, CDCl3) $\delta$ 177.03, 163.37, 160.93, 134.51, 134.48, 129.40, 129.32, 115.60, 115.39, 47.55, 42.67, 31.73, 28.10, 26.61; LCMS: m/z 249.78 [M+H]+.

##### S1.2.15 Experimental Settings for ADMET Bench Testing

**Cytotoxicity (HepG2) Assay** Cytotoxicity was evaluated using the human hepatocellular carcinoma cell line HepG2 (ATCC). Cells were maintained in ATCC-formulated Eagle's Minimum Essential Medium (EMEM) supplemented with 2% heat-inactivated fetal bovine serum and 1× penicillin–streptomycin. For assays, cells were grown to 80–100% confluency, detached using TrypLE, and resuspended in assay medium. Cell viability was confirmed to be $\geq$ 90% by trypan blue exclusion prior to seeding. HepG2 cells were dispensed into white solid-bottom 1536-well plates at a density of 50,000 cells/mL (5 $\mu$L per well; about 250 cells/well) using a MultiFlo dispenser and incubated at 37 °C with 5% $CO_2$ for 72 hours. Following incubation, plates were equilibrated to room temperature and cell viability was assessed by addition of CellTiter-Glo reagent (2 $\mu$L per well). Luminescence was measured after a 10 min incubation using a ClarioStar plate reader with a 1536-well aperture. Luminescent signal was used as a measure of ATP content and normalized to untreated controls to determine relative cell viability.

**HDF Cytotox Assay** Human dermal fibroblasts were cultured with Dulbecco's modified eagle medium supplemented with 10% fetal bovine serum, 100units/mL of Penicillin, and 100$\mu$g/mL of Streptomycin in 5% $CO_2$ incubator at 37 °C. Cells were incubated until a confluency of 70% was obtained and then detached from the flask with trypsinization. To avoid cell death, once cells were detached, fetal bovine serum-supplemented media was added. Cells were centrifuged, counted, and then diluted to a cell concentration of 64,000 cells/mL. In a 384-well plate, 49$\mu$L of the cell suspension was added to each well. Compounds were serially diluted and 1$\mu$L was added to respective wells starting at a final assay compound concentration of 100$\mu$M. The plate was incubated for 3 days in a 5% $CO_2$ incubator at 37°C. After incubation, Resazurin was added, and fluorescence was read after 18 hours of incubation.

**Liver Microsome Assay** Compounds were plated at a final concentration of 1$\mu$M into a 25mM Potassium Phosphate Buffer (pH 8) containing 0.5mg/mL of human liver microsomes and 2mM

$MgCl_2$. The plate was incubated and shaken for 10 min. at 37 °C and 100rpm. Immediately after incubation, NADPH at an assay concentration of 1mM was added to begin the reaction. At different time points, 50$\mu$L of the reaction mix was pipetted into a plate containing 250$\mu$L of acetonitrile with 0.1% Formic Acid to stop the reaction. The plate was then shaken for another 15 minutes at 800rpm at room temperature and centrifuged for 25 minutes at 3700rpm. Samples were then run in a Bruker QTOF III mass spec to obtain data. Each drug was run in triplicate with a standard curve to quantify concentration.

**Kinetic Solubility Assay** A DMSO stock solution of each compound was diluted to a final compound concentration of 100$\mu$M into wells containing 0.1M Sodium Phosphate Buffer (pH 7.4). A clean stir bar was added to each well, and the solutions were stirred for 24 hours at a stirring speed of 150rpm at room temperature. After 24 hours, the plate was centrifuged at 1000 G for 2 minutes. The supernatant was diluted to a theoretical compound concentration of 10$\mu$M in more of the same buffer. These samples were run and analyzed by LC-MS along with a 10$\mu$M solution of each compound in methanol. Using AUC values, each assayed solution was compared to its methanol drug solution counterpart to obtain the solubility percentage.

**Plasma Protein Binding Assay** Each compound was mixed into plasma at a concentration of 10$\mu$g/mL. The mixtures were incubated at room temperature for 15 minutes. Using a Thermoscientific "Rapid Equilibrium Dialysis" plate, 300$\mu$L of PBS and 100$\mu$L of plasma were plated into the white and red wells respectively. The plate was shaken at 250 rpm in an incubator for 5 hours at a temperature of 37 °C. After incubation, 50$\mu$L of fresh plasma was mixed with 50$\mu$L of the PBS samples, and 50$\mu$L of fresh PBS was mixed with 50$\mu$L of the plasma samples in centrifuge tubes. 200$\mu$L of water, 20$\mu$L of water spiked with 1 $\mu$g/mL of Warfarin, and 180$\mu$L of Acetonitrile were added to each sample. Samples were vortexed and centrifuged at 14000 rpm for 5 minutes and run in LCMS where % bound was calculated based on the mass intensities in comparison to warfarin AUC values.

#### S1.3 Fleming Prompts

**Fleming Operation and Tool Prompts** Prompts used in the Fleming operation and tools can be found on our GitHub repository.

**Plausibility Assessment** You are an experienced biochemist looking for novel M.tb antibiotics. I'll provide you with a SMILES structure. Please analyze the SMILES structure in detail and let me know if you think this molecule is worth bench-testing for Mtb antibiotics. Keep in mind that bench testing is expensive. Respond only with a decision between Possible/Not-Likely with a concise reasoning. Respond in the format: Possible/Not-Likely, Reasoning. SMILES Structure: {smiles}

**Fleming Benchmarking - Fleming** You are an expert medicinal chemist. You will be presented with a molecule that has secret data, it is a great clinical candidate as a drug against TB, or a random drug-like molecule. You will need to report the output probability of the molecule as the drug against TB. Report the probability as a number between 0 and 1, where 1 is that you are certain that it would be a good drug against TB, and 0 is that you are certain that it is not a valid drug against TB. These may be novel molecules, drugs in development, or repurposed drugs that have then been tested against TB. Return the probability as a number between 0 and 1. Return just the number. Here is the smiles: {smiles}. Please do a full literature review, not the brief literature review, and also use the inhibition model to help guide your prediction.

**Fleming Benchmarking - GPT o4-mini** You are an expert medicinal chemist. You will be presented with a molecule that has secret data that it's a great clinical candidate as a drug against TB, or a random drug-like molecule. You will need to report the output probability of the molecule as the drug against TB. Report the probability as a number between 0 and 1, where 1 is that you are certain that it would be a good drug against TB, and 0 is that you are certain that it is not a valid drug against TB. These may be novel molecules, drugs in development, or repurposed drugs that have then been tested against TB. Return the probability as a number between 0 and 1. Return just the number. Here is the smiles: {smiles}. Please use your knowledge base to find the answer.

### S2 Supplementary Figures

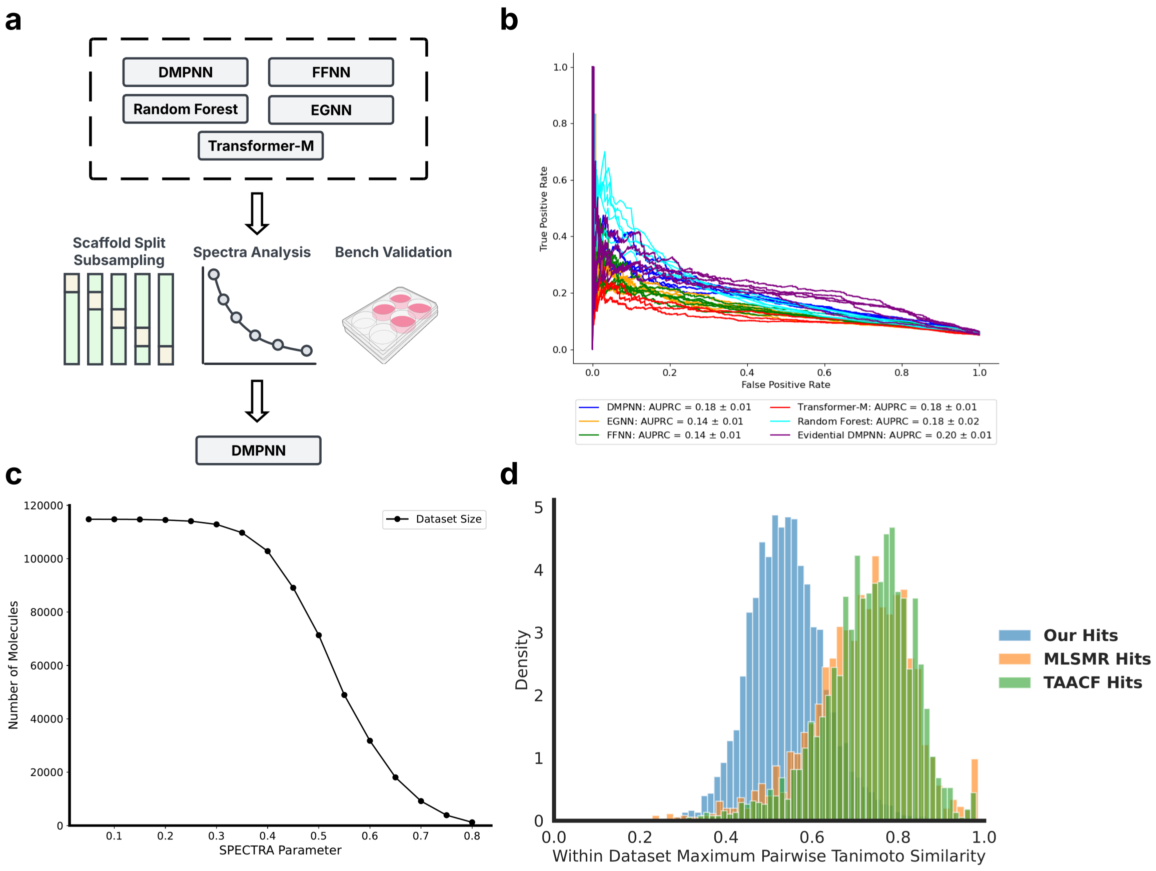


**Supplementary Figure 1: Inhibition Prediction Model Performance (a)** Illustration of the benchmarking methods we applied to determine the best model architecture for Mtb growth inhibition prediction. **(b)** Area under Precision-Recall Curve (AUPRC) of benchmarked models for Mtb growth inhibition prediction. **(c)** The size of the dataset as the SPECTRA[1] parameter increases (i.e., the model contains more diverse molecules) **(d)** The maximum pairwise Tanimoto similarity of hits within the MLSMR, and our dataset.

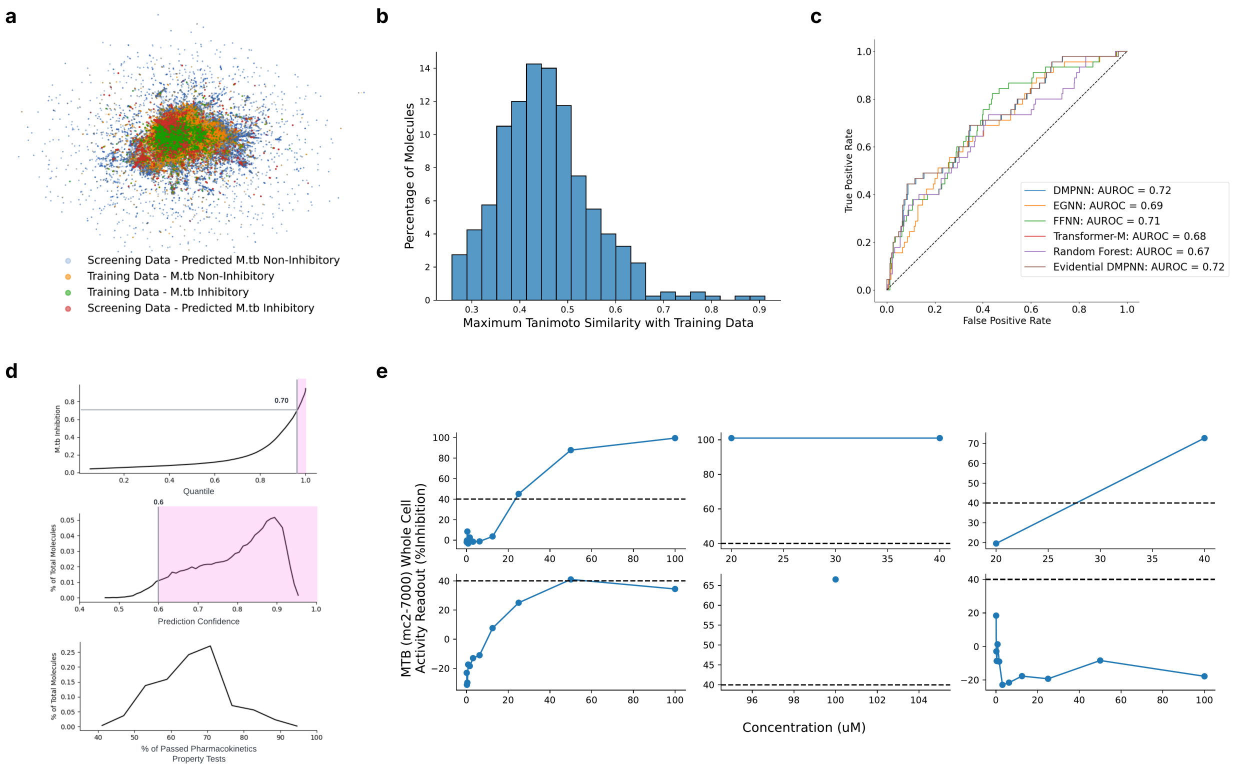


**Supplementary Figure 2: *In-vitro* Library Screening Validation (a)** Chemical space illustration of the training dataset and the screening library using UMAP on Morgan fingerprints of the molecules. **(b)** The distribution of maximum pairwise Tanimoto similarity of the screening library and the training dataset. **(c)** Area under Receiver Operating Characteristic (AUROC) curve of the benchmarked models on the *In-vitro* screening dataset. **(d)** The predicted Mtb growth inhibition and the prediction confidence using Evidential DMPNN, as well as the predicted ADMET profile using ADMET.AI models of the screening dataset. **(e)** Dose-response curve of the six predicted inhibitory molecules in the screening dataset using Evidential DMPNN.

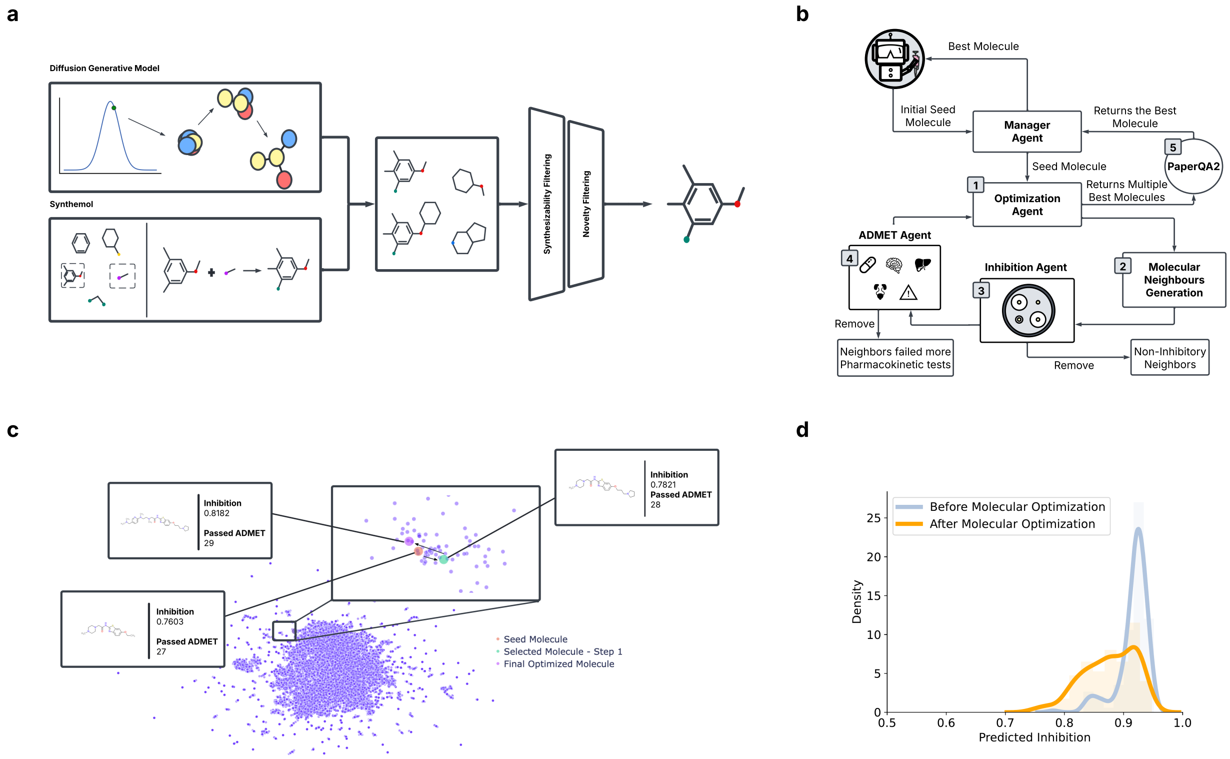


**Supplementary Figure 3: Molecular Generation and Optimization (a)** Illustration of Molecular Generation agent, where Fleming generates Mtb inhibitory molecules from a diffusion model and a SyntheMol model. **(b)** Illustration of Molecular Optimization agent, where the molecules are optimized towards the desired property while maintaining it's predicted Mtb growth inhibition. **(c)** Example of molecular optimization where the initial molecule, which only passes 27 TDC ADMET properties (orange), undergoes two steps of optimization (orange to green to purple), and now has passed 29 TDC ADMET properties. **(d)** Distribution of predicted Mtb growth inhibition before and after molecular optimization. After optimization, inhibition probability was stable (median probability 0.92 to 0.88)

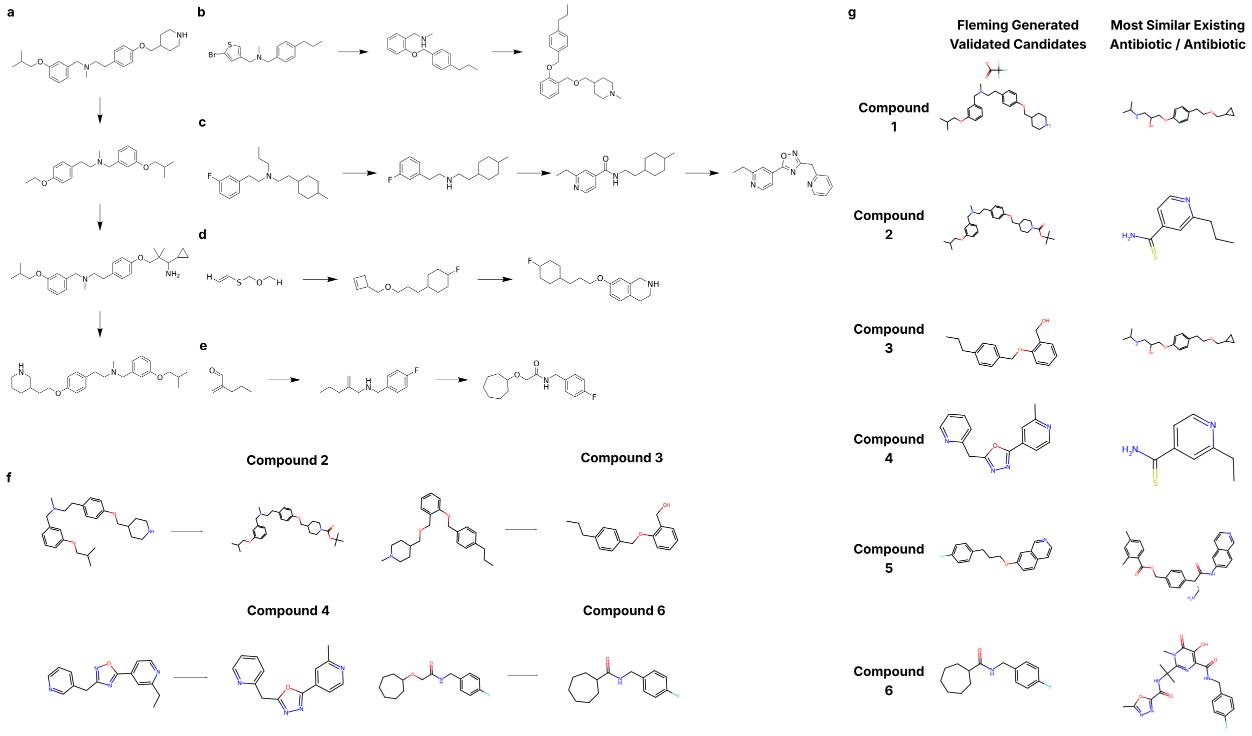


**Supplementary Figure 4: *In-vitro* Bench Validation of Fleming Designed Molecules (a-e)** The steps taken in the Molecular Optimization agent for all 6 Fleming-designed molecules. **(f)** The minor modifications made to the Fleming-designed molecules by medicinal chemists. **(g)** The most similar existing antibiotics compared to the Fleming-designed molecules using Tanimoto similarity.

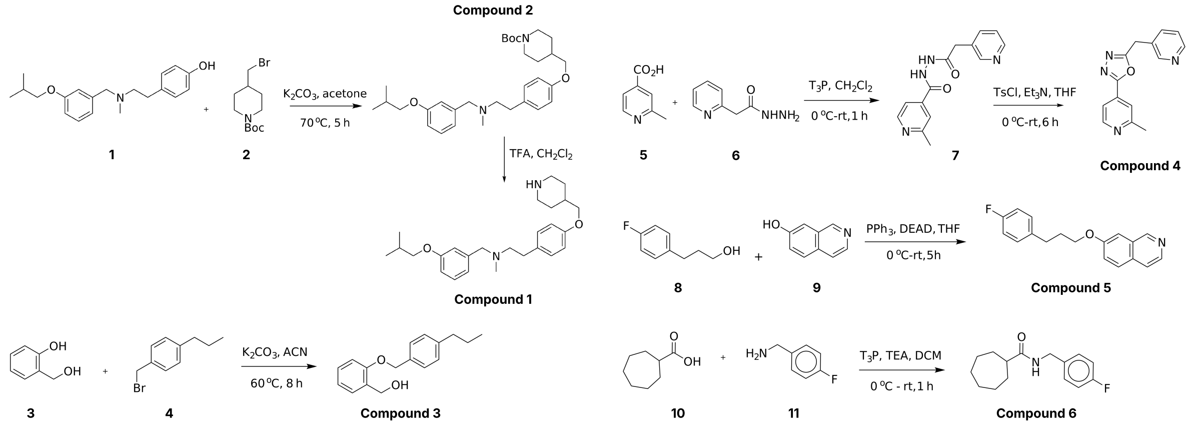


**Supplementary Figure 5: Synthesize pathways of the six Fleming designed molecules**

### S3 Supplementary Tables

*** Supplementary Table 3-6 are in the attached Excel sheet due to the size of the tables.**

**Supplementary Table 1:** Fleming Tool Description Table

| Tool Name | Tool Description |
|---|---|
| Inhibition Prediction Model | Predicts MTB growth inhibition (with prediction confidence) |
| MCTS Rationale Extraction Model | Using MCTS [15] to extract rationales for activity |
| ADMET.AI Prediction Models | Run ADMET.AI[10] models and binarized into pass/fail + score or return detailed numerical predictions |
| Cytotoxicity Prediction Model | Run cytotoxicity prediction model built using data from [7] |
| BBBP Prediction Model | Run Blood-brain barrier penetration prediction using dataset from MoleculeNet [13] |
| Side Effect Prediction Model | Run using the datasets from MoleculeNet [13] |
| Molecule Generation Model - Diffusion | Generate MTB inhibitory molecules from the diffusion model |
| Molecule Generation Model - SyntheMol | Generate MTB inhibitory molecules from SyntheMol [15] |
| Calculate Novelty and Synthesizability | Calculate molecule similarity and synthetic accessibility (SAScore[20] and SCScore[21]) |
| Generate Molecular Neighbors | Find chemical neighbors prioritized for ADMET and Inhibition using Synspace[26] |
| Get IUPAC Name | Get IUPAC name for a molecule |
| Get Molecule Image | Visualize the molecule using RDKit |
| Get RDKit Documentation | Retrieve RDKit documentation for running cheminformatic tasks |
| Check Drug Likeness | Check if the molecule fails any drug-likeness filters |
| Molecule Describer | Describes the structure of the molecule in natural language |
| Quick GPT-based Literature Review | Literature search using GPT |
| PaperQA2 Molecule Analysis | Time-intensive substructure-based literature search using PaperQA2[27] |
| Request for User Clarification | An interactive tool to request clarification from the user |
| Run Python Scripts | Executes Python for custom analysis |

**Supplementary Table 2:** Pharmacokinetics pass/fail thresholds determined by existing MTB antibiotics used in ADMET Agent

| Higher the Better | | Lower the Better | | | |
|---|---|---|---|---|---|
| Task | Lower Bound | Task | Higher Bound | Task | Higher Bound |
| Bioavailability | 0.2449 | Mutagenicity (AMES) | 0.7347 | Drug Induced Liver Injury (DILI) | 0.4365 |
| Human Intestinal Absorption | 0.0310 | Blood Brain Barrier Penetration | 0.7612 | NR-AR-LBD | 0.0864 |
| Solubility | -4.147 | CYP1A2 Inhibition | 0.0574 | NR-AR | 0.1867 |
| Cell Effective Permeability | -6.922 | CYP2C19 Inhibtion | 0.1286 | NR-AhR | 0.1559 |
| PAMPA Permeability | 0.0217 | Plasma Protein Binding Rate (PPBR) | 89.15 | NR-Aromatase | 0.1304 |
| Hepatocyte Clearance | -6.7 | CYP2C9 Inhibition | 0.10 | NR-ER-LBD | 0.07978 |
| Microsome Clearance | 11.57 | CYP2D6 Inhibition | 0.0457 | NR-ER | 0.2302 |
| Half-Life | -13.92 | CYP3A4 Inhibition | 0.2720 | NR-PPAR-gamma | 0.0365 |
| Volume of Distribution at Steady State (VDss) | -4.141 | Carcinogenicity | 0.3685 | SR-ATAD5 | 0.0733 |
| | | Clinical Toxicity | 0.4365 | SR-ARE | 0.3770 |
| | | SR-MMP | 0.3877 | SR-HSE | 0.0646 |
| | | SR-p53 | 0.2093 | Skin Reaction | 0.6774 |
| | | hERG | 0.5476 | P-glycoprotein Inhibition | 0.7790 |
| | | Acute Toxicity LD50 | 2.8784 | Lipophilicity | 1.4683 |
